## Supplementary Figures and Tables for "A unified model for interpretable latent embedding of multi-sample, multi-condition single-cell data"

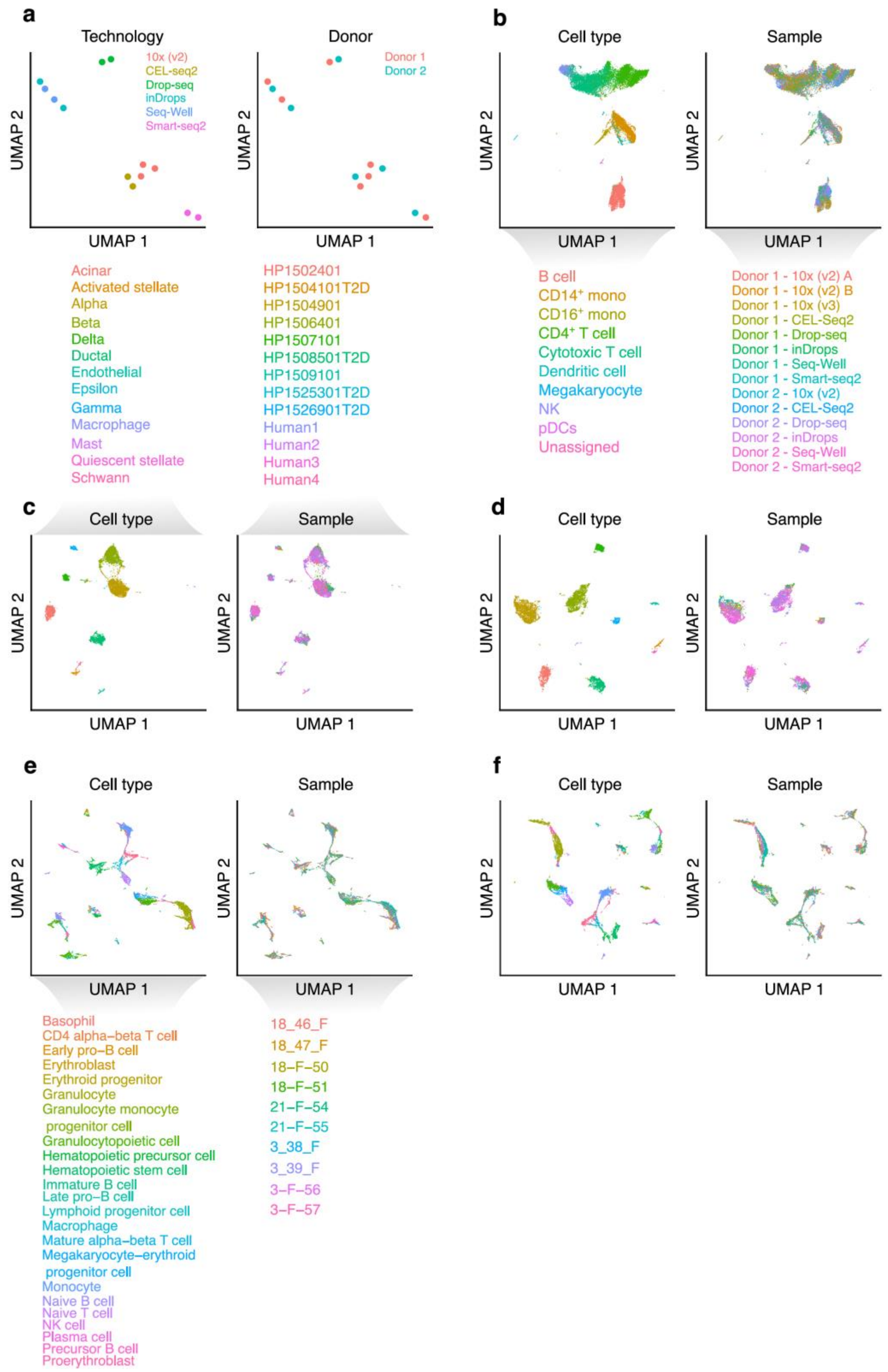

**Supplementary Figure 1.** Analysis of sample-to-sample variability with GEDI
(Continued on the next page)

(**a**) UMAP embedding of the sample-specific manifold distortions learned by GEDI for the PBMC dataset[1] (related to **Fig. 2a**). For each sample *i*, GEDI learns a set of sample-specific manifold parameters, consisting of $\Delta\boldsymbol{Z}_i$ and $\Delta\boldsymbol{o}_i$ (see **Methods** for the explanation of these parameters). We concatenated $\Delta\boldsymbol{Z}_i$, followed by vectorization, to obtain a vector $\boldsymbol{\theta}_i \in \mathbb{R}^{G(K)}$ for each sample *i*, where $G$ is the number of genes and $K$ is the number of principal axes (see **Methods**). We then regressed out the effect of donor from each element $j$ of each $\boldsymbol{\theta}_i$ by modeling $\theta_{i,j} \sim \alpha_j + \beta_j h_i$ across the samples and taking the residual of the regression (here, $h_i$ represents the donor for sample $i$). We then performed PCA and UMAP dimensionality reduction on the residuals to obtain the plots shown here. Each dot represents one sample, labeled by the single-cell technology used (left) or donor of origin (right). Only technologies with more than one sample are displayed. A similar analysis was performed to obtain **Fig. 2a**, with the difference that in that figure, the effect of technology was regressed out ($h_i$ was set to the technology used for each sample $i$) (**b**) UMAP embedding of the cells in the PBMC dataset after integration with GEDI (hyperplane mode). Each dot represents one cell, colored by the cell type labels from the original study (left) or by sample (right). This figure is related to **Fig. 2b**, with the difference that **Fig 2b** is based on GEDI in the hyperellipsoid mode. (**c**) UMAP embedding of the cells in the Pancreas dataset[2, 3] after integration with GEDI (hyperellipsoid mode). Each dot represents one cell, colored by the cell type labels from the original study (left) or by sample (right). (**d**) Same as (c), but integration was performed with GEDI (hyperplane mode). (**e**) UMAP embedding of the cells in the Tabula Muris dataset[4] after integration with GEDI (hyperellipsoid mode). Each dot represents one cell, colored by the cell type labels from the original study (left) or by sample (right). (**f**) Same as (e), but integration was performed with GEDI (hyperplane mode).

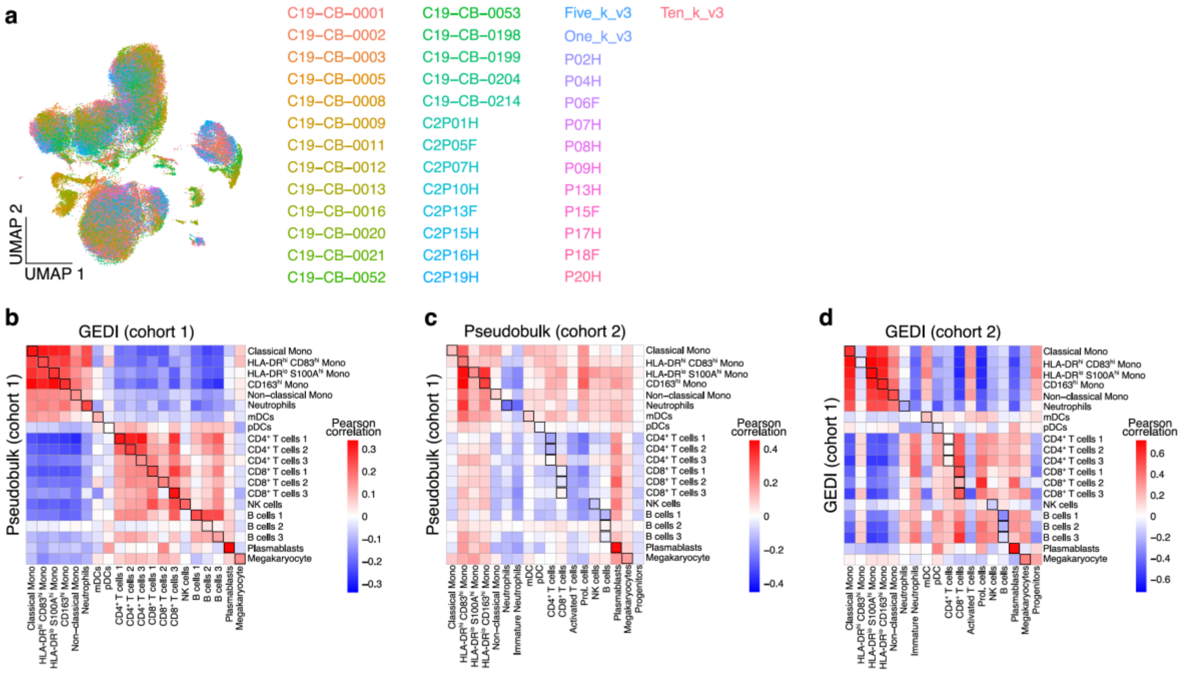

**Supplementary Figure 2.** Additional comparisons between GEDI and pseudo-bulk analysis
(**a**) UMAP embedding of the COVID-19 dataset[5] for cohort 1 (similar to **Fig. 3b**). The color indicates the donor labels from the original study. (**b**) Comparison between the mean transcriptomic vector per cell type, obtained from GEDI, and differential gene expression values (log fold-change) obtained from pseudo-bulk analysis, for the comparison of severe COVID-19 vs. control cases in cohort 1 (related to **Fig. 3e**). Heatmap shows the Pearson correlation values between GEDI (columns) and pseudo-bulk analysis (rows). (**c**) Same as in (b) but showing reproducibility between cohort 1 (rows) and cohort 2 (columns) for the pseudo-bulk analysis (related to **Fig. 3f**). (**d**) Same as in (b-c) but showing reproducibility between cohort 1 (rows) and cohort 2 (columns) for GEDI (related to **Fig. 3g**).

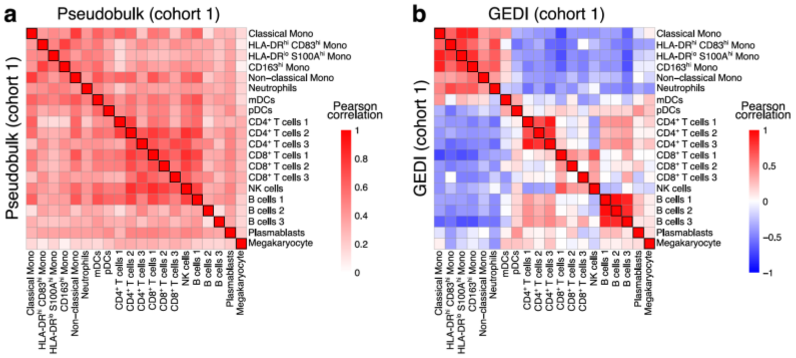

**Supplementary Figure 3.** GEDI differential expression estimates preserve cell type-specificity
(**a**) Pseudo-bulk differential expression (DE) analysis between mild COVID-19 vs. control in cohort 1. Heatmap shows the Pearson correlation for the estimated DE profiles of different cell types. (**b**) same as (a) but showing the Pearson correlation of the mean transcriptomic vectors of the cell types, obtained from GEDI.

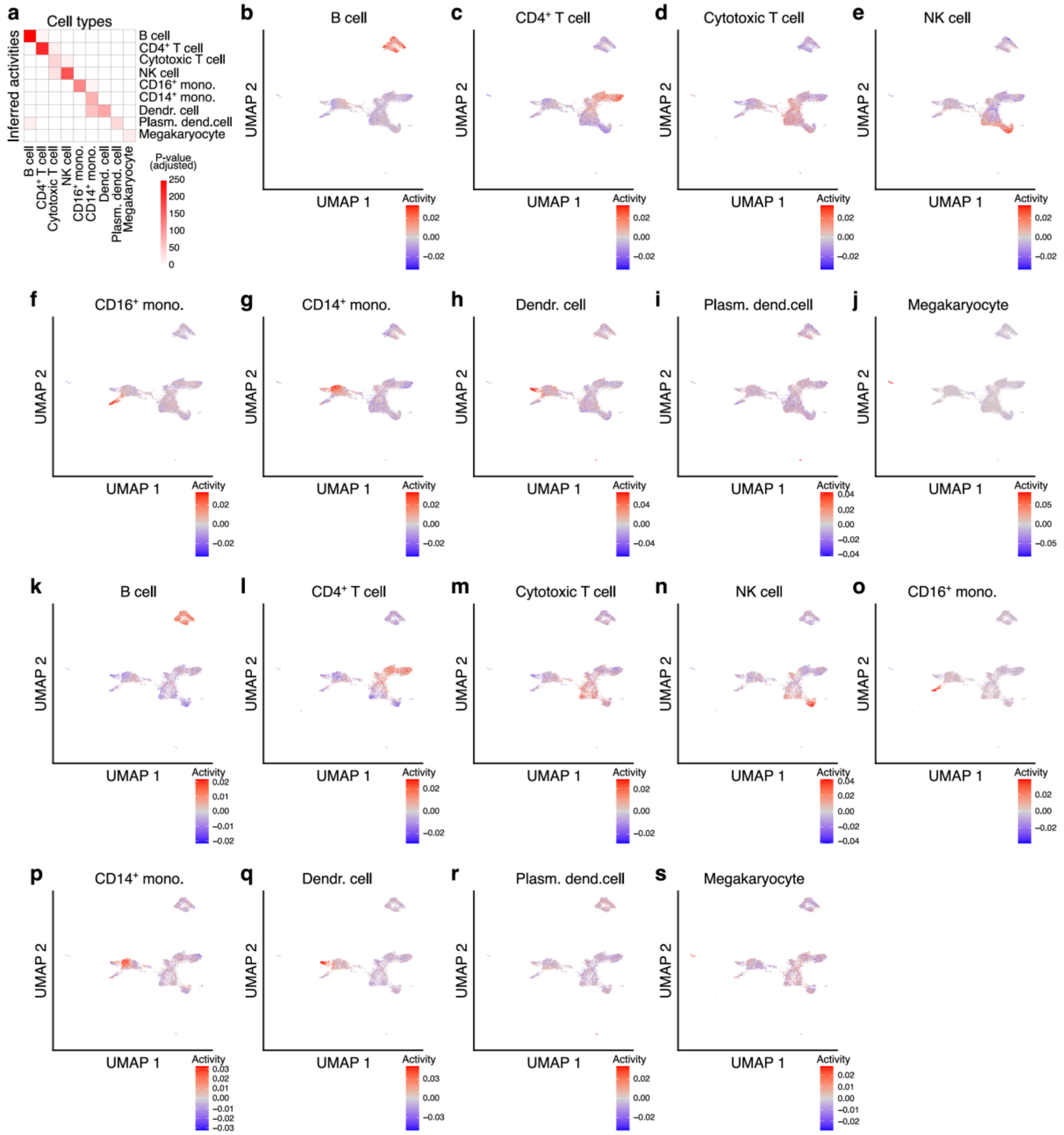

**Supplementary Figure 4.** Projection of cell type signatures with GEDI
(**a**) Cell type signature projections obtained by GEDI are compared to the true labels in the PBMC dataset (donor 2). Heatmap shows adjusted p-values for differential enrichment of inferred cell type signatures from GEDI (rows) for each cell type (columns). This figure is similar to **Fig. 4a**, with the difference that, here, the gene signatures are obtained from donor 1, followed by their projection on the cells from donor 2. (**b-j**) UMAP plots showing single-cell projection of cell-type signature activities for donor 1. (**k-s**) UMAP plots showing single-cell projection of cell-type signature activities for donor 2.

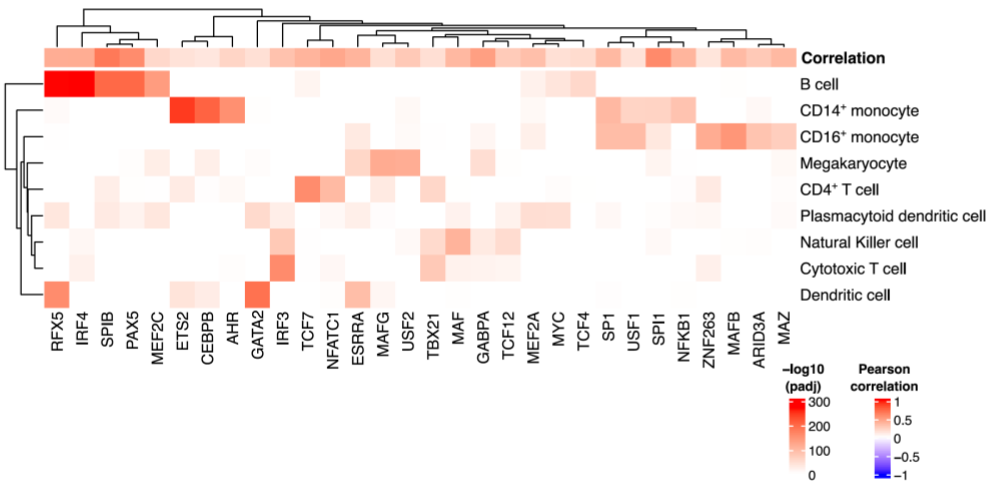

**Supplementary Figure 5.** Identification of cell type-specific transcription factor activities with GEDI
Heatmap shows adjusted p-values for differential enrichment of inferred transcription factor (TF) activities from GEDI (columns) for each cell type (rows) in the PBMC dataset. The top row shows the Pearson correlation between inferred activity and model-fitted mRNA abundance. For this figure, we included only TFs with Pearson correlation >0.1 between activity and model-fitted mRNA abundance. Then, we show only the top 30 most significant TFs based on sorting the minimum p-value per TF.

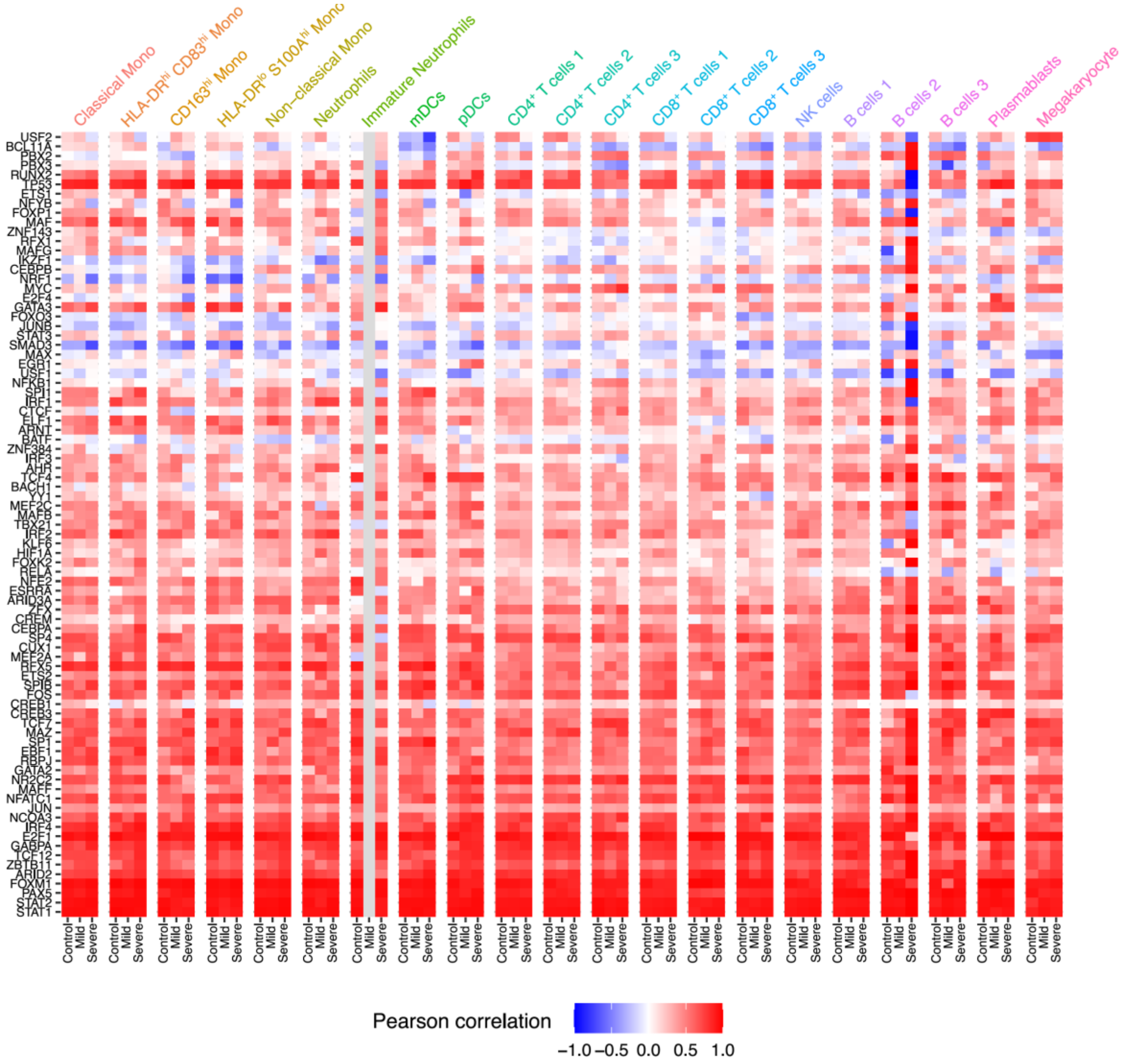

**Supplementary Figure 6.** Comparison between the projected TF regulon activities and TF mRNA abundance
The heatmap shows the Pearson correlation between the inferred regulon activity of each TF and the GEDI-imputed abundance of the mRNA encoding that TF, separately calculated for each cell type and each COVID-19 condition in the COVID-19 cohort 1 dataset.

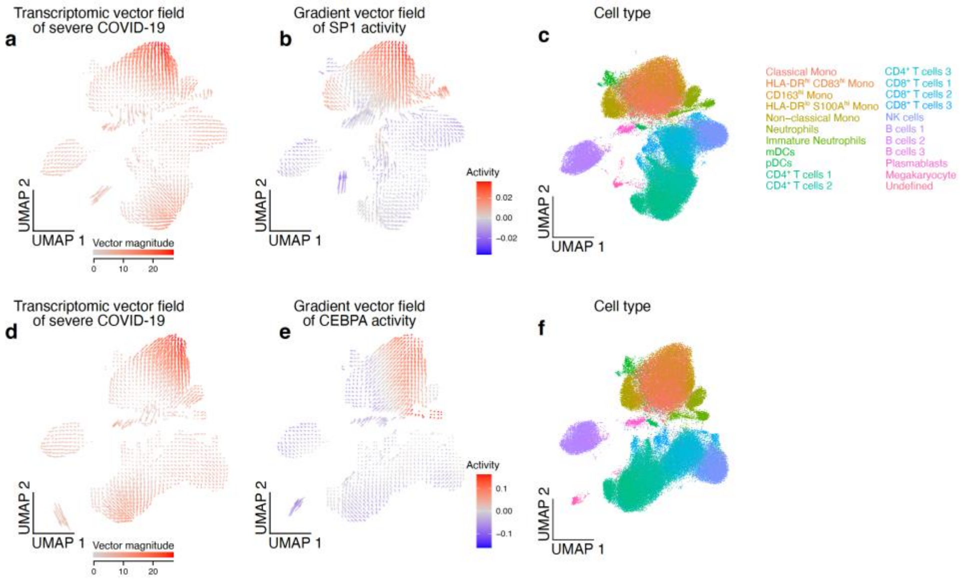

**Supplementary Figure 7.** Correlation of TF activity gradient vectors and the transcriptomic vector field
(**a-c**) The activity gradient of SP1 correlates with the transcriptomic vector of severe COVID-19 in monocytes. (**a**) UMAP representation of the transcriptomic vector field of severe COVID-19. The color shows the vector magnitude. (**b**) Gradient vector field of SP1 activity. The color represents SP1 activity. (**c**) The same UMAP as in (a-b), but the color represents the cell type labels as a reference. (**d-f**) same as in (a-c), but for CEBPA. (**d**) Same as (a), but UMAP coordinates are derived from the analysis of the vector field of CEBPA. (**e**) Same as (b) but showing CEBPA activity. (**f**) Same as (c), but UMAP coordinates are derived from the analysis of the vector field of CEBPA. Also see **Fig. 4d-g**.

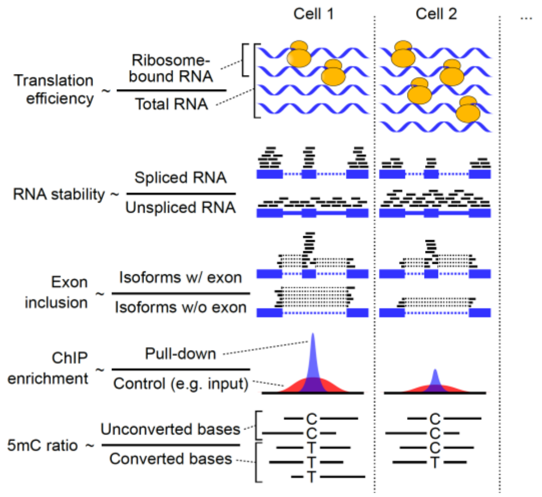

**Supplementary Figure 8.** Examples modalities that are defined as the ratio of two quantities.

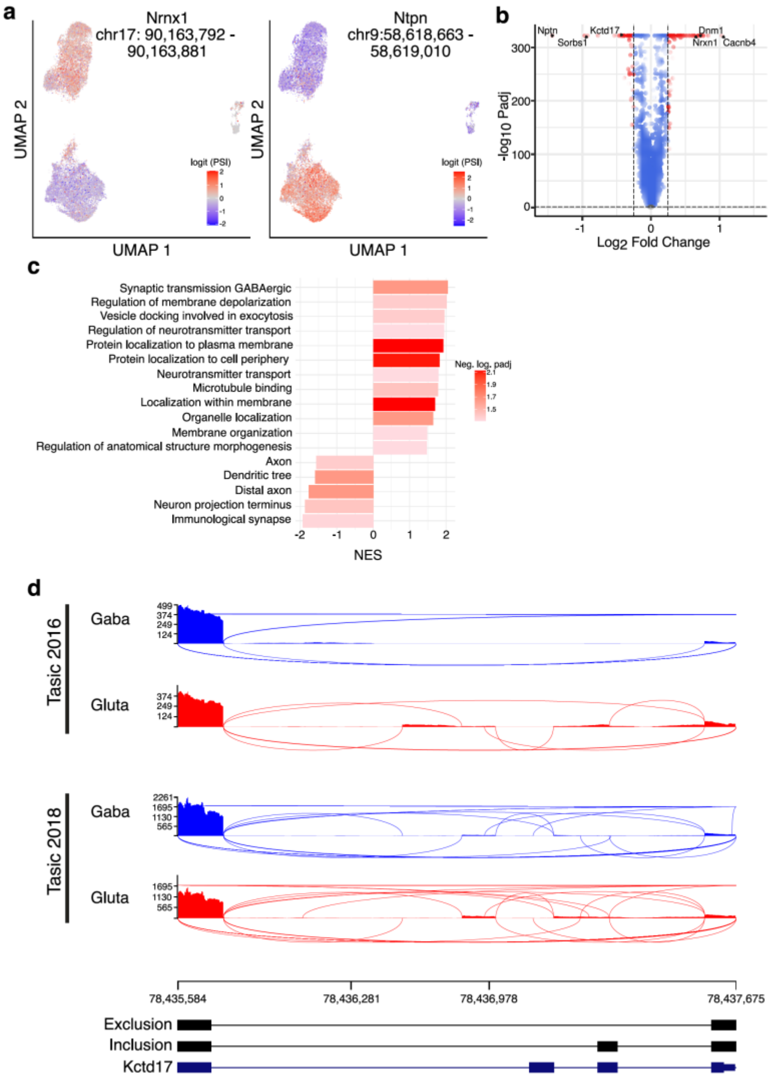

**Supplementary Figure 9.** Analysis of the latent splicing space learned by GEDI

(**a**) UMAP embedding of the latent splicing space of mouse cortical cells after integration of data from two studies[6, 7]. This figure is similar to **Fig. 5d**, with the difference that the color represents the log-odds of inclusion/exclusion for a cassette exon in Nrnx1 (left) and a cassete exon in Ntpn (right). (**b**) Volcano plot showing differential PSI between GABAergic and Glutamatergic neurons. Red points indicate exon inclusion events with a $\log_2$ fold-change >0.25. (**c**) Gene Set Enrichment Analysis (GSEA[8]) of differential exon inclusion between GABAergic and Glutamatergic neurons. Bar height indicates the normalized enriched score while bar color represents the negative $\log_{10}$ of the FDR-adjusted p-value. (**d**) Sashimi plots of an example cassette exon in Kctd17 that is differentially spliced between neuronal subtypes.

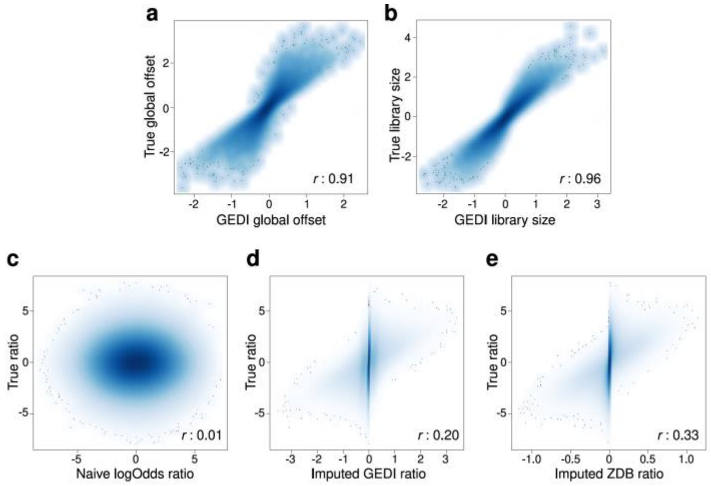

**Supplementary Figure 10.** Analysis of simulated pairs of UMI counts with GEDI

To simulate paired UMI counts (similar to, for example, UMI counts that are observed from analysis of spliced and unspliced RNA), we started with matrix $\boldsymbol{M}$ containing UMI counts for 11,532 genes across 17,180 cells from a real scRNA-seq dataset (PBMC dataset[1] donor 1). We used the matrix $\boldsymbol{M}$ to create two other matrices of the same dimensions, $\boldsymbol{M}_1$ and $\boldsymbol{M}_2$, based on binomial subsampling of each element $m_{g,n}$ of $\boldsymbol{M}$ according to the distribution $m_{1,g,n}$~B($m_{g,n}$,$p_{g,n}$), and $m_{2,g,n}$= $m_{g,n}$– $m_{1,g,n}$. The sampling probability $p_{g,n}$ for each gene $g$ in each cell $n$ was chosen as $p_{g,n} = 1/\left[1 - \exp\left[-(\boldsymbol{ZB})_{g,n} + o_g + s_n\right]\right]$, where $\boldsymbol{Z}$ and $\boldsymbol{B}$ are two matrices of rank 20, with each element sampled from $N(0,0.5)$, and $\boldsymbol{o}$ and $\boldsymbol{s}$ are vectors representing gene-specific and cell-specific offsets, respectively, sampled from $N(0,1)$. (**a**) Comparison of the ground-truth gene-specific offset $\boldsymbol{o}$ with GEDI-inferred gene-specific offsets. Pearson correlation is shown. (**b**) Comparison of the ground-truth cell-specific offset $\boldsymbol{s}$ with GEDI-inferred cell-specific offsets. (**c**) Comparison of the ground-truth logit of $p_{g,n}$ to a naïve estimator, obtained as $log[(m_{1,g,n}+1)/ (m_{2,g,n}+1)]$. Each data point represents one gene in one cell. (**d**) Comparison of the ground-truth logit of $p_{g,n}$ to the GEDI-imputed values, i.e., the expected value of the latent $y_{g,n}$ given the observed (simulated) $m_{1,g,n}$ and $m_{1,g,n}$ UMI counts and the model-predicted value. (**e**) Comparison of the ground-truth logit of $p_{g,n}$ to the model-predicted values.

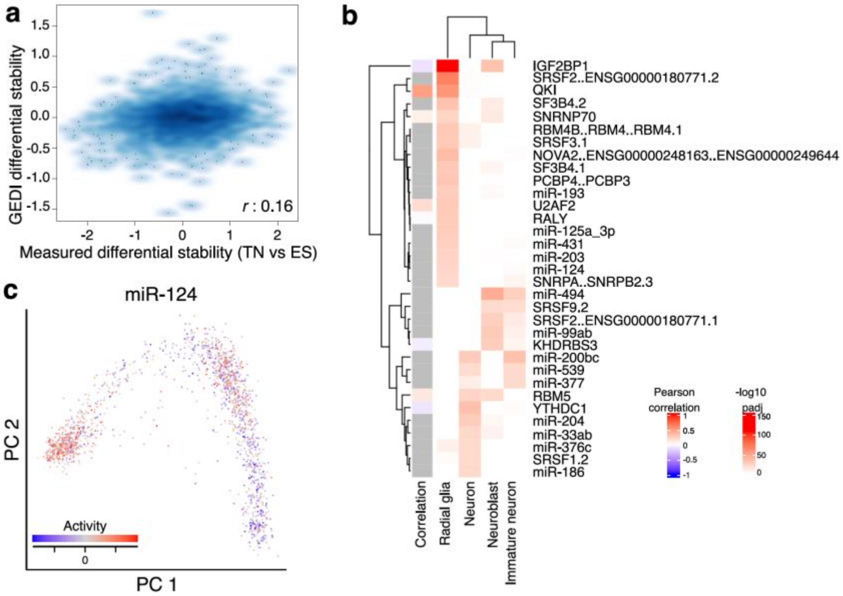

**Supplementary Figure 11.** Analysis of the latent stability space learned by GEDI

(**a**) Comparison between differential stability estimates inferred from GEDI vs. RNA half-life measurements obtained from mouse embryonic stem cells (ESCs) and in vitro-differentiated terminal neurons (TNs)[9]. GEDI estimates were obtained after analyzing the ratio of spliced and unspliced transcripts at the single-cell level in a model of sensory neurogenesis[10]. Differential stability estimates were obtained by calculating the slope of the imputed log-ratio of spliced vs. unspliced transcripts vs. pseudotime, which represents stability changes per unit of time. Pseudotime scores were obtained from the original publication[10]. Pearson correlation between GEDI estimates and experimental measurements of stability is shown. (**b**) GEDI identifies cell type-specific activities of post-transcriptional regulators. GEDI was applied to analyze the ratio of unspliced and spliced RNAs in human neurons, using a previously published dataset of human embryonic glutamatergic neurogenesis[11]. In this analysis, we modeled the spliced/unspliced latent manifold as a function of the regulatory networks of RNA binding proteins (RBPs) and miRNAs. Heatmap shows adjusted p-values for differential enrichment of inferred post-transcriptional regulator activities from GEDI (rows) for each cell type (columns). Left annotation heatmap shows Pearson correlation between inferred activity and model-fitted mRNA abundance (for RBPs). Gray values indicate not available values of expression of miRNAs. (**c**) PCA of the human embryonic glutamatergic dataset[11]. The color shows the projected regulon activity of miR-124, which is an inhibitor of mRNA stability[12]. Activity pattern shows that the targets of miR-124 are more stable in radial glia and less stable in differentiated neurons, consistent with higher miR-124 activity in differentiated neurons.

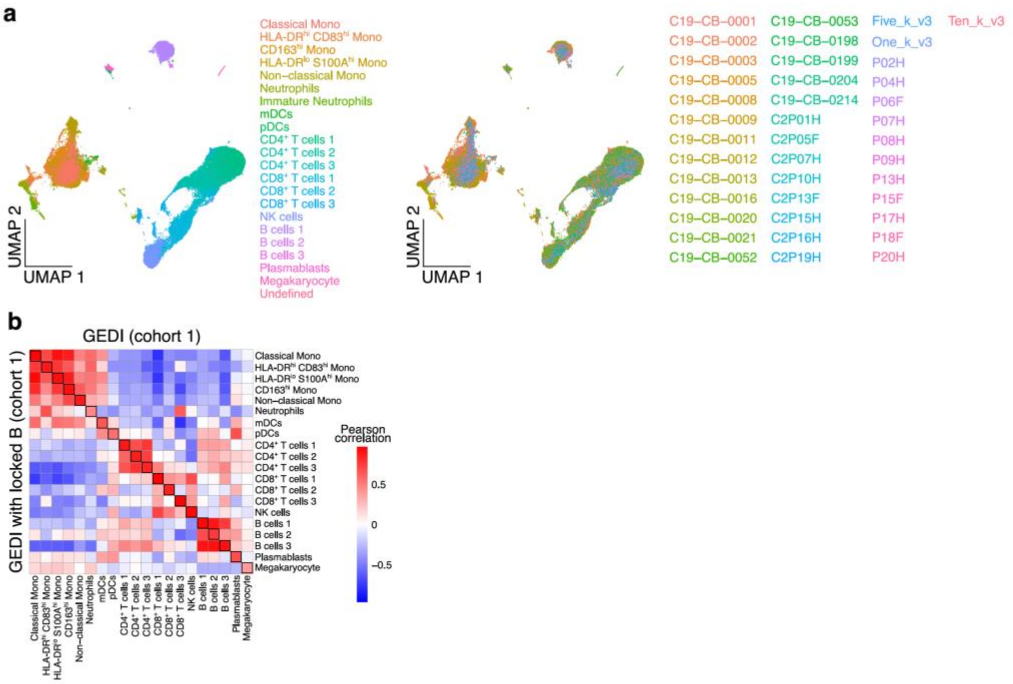

**Supplementary Figure 12.** Post-hoc analysis of prespecified integrated cell state spaces with GEDI
(**a**) UMAP embedding of the cells in the COVID-19 cohort 1 dataset after integration with Harmony[13]. Each dot represents one cell, colored by the cell type labels from the original study (left) or by sample (right). (**b**) The mean transcriptomic vector field, representing expression shift in mild COVID-19 relative to healthy controls, was obtained using GEDI based on the integrated cell state space from Harmony, or by *de novo* integration of data with GEDI. The heatmap shows the pairwise Pearson correlations of the mean transcriptomic vectors of cell types between the Harmony-based GEDI analysis (rows) and *de novo* GEDI analysis (columns).

**Supplementary Table 1.** List of source data files

The Zenodo records are available at https://zenodo.org/record/8222040 (DOI: 10.5281/zenodo.8222040) and https://zenodo.org/record/8222698 (DOI: 10.5281/zenodo.8222698).

| |
|---|
| **File name:** pbmc_lis_K.rds (related to Figure 2)<br>**Description:** List with embeddings for the integration of PBMC data across various number of latent variables (*K*) and integration methods. For each *K* and method, three objects are included: ‘embedding_res’: embeddings of the integrated method; ‘umap_2_res’: UMAP embeddings with two dimensions; ‘umap_n_res’ UMAP embeddings with *K* dimensions.<br>**Download URL:** https://zenodo.org/record/8222040/files/pbmc_lis_K.rds?download=1 |
| **File name:** pancreas_lis_K.rds (related to Figure 2)<br>**Description:** List with embeddings for the integration of Pancreas data. Similar to above.<br>**Download URL:** https://zenodo.org/record/8222040/files/pancreas_lis_K.rds?download=1 |
| **File name:** tabulaMuris_lis_K.rds (related to Figure 2)<br>**Description:** List with embeddings for the integration of Tabula Muris data. Similar to above.<br>**Download URL:** https://zenodo.org/record/8222040/files/tabulaMuris_lis_K.rds?download=1 |
| **File name:** pbmc_gedi_model_bothDonors.rds (related to Figure 2)<br>**Description:** GEDI object of PBMC data.<br>**Download URL:** https://zenodo.org/record/8222040/files/pbmc_gedi_model_bothDonors.rds?download=1 |
| **File name:** COVID19_gedi_model_bothCohorts.rds (related to Figure 2)<br>**Description:** GEDI object of COVID-19 data (two cohorts).<br>**Download URL:** https://zenodo.org/record/8222040/files/COVID19_gedi_model_bothCohorts.rds?download=1 |
| **File name:** COVID19_gedi_model_cohort1.rds (related to Figure 3)<br>**Description:** GEDI object of COVID-19 data (cohort1), with sample-level variables incorporated in the model.<br>**Download URL:** https://zenodo.org/record/8222040/files/COVID19_gedi_model_cohort1.rds?download=1 |
| **File name:** COVID19_gedi_model_cohort2.rds (related to Figure 3)<br>**Description:** GEDI object for COVID-19 data (cohort2), with sample-level variables incorporated in the model.<br>**Download URL:** https://zenodo.org/record/8222698/files/COVID19_gedi_model_cohort2.rds?download=1 |
| **File name:** COVID19_list_DE.rds (related to Figure 3)<br>**Description:** List with Differential Expression results for DESeq2 and GEDI, for the "severe_vs_control" and "mild_vs_control" comparisons.<br>**Download URL:** https://zenodo.org/record/8222698/files/COVID19_list_DE.rds?download=1 |
| **File name:** pbmc_gedi_model_donor1_celltype.rds (related to Figure 4)<br>**Description:** GEDI object of PBMC data (donor 1), with prior information of cell type signatures incorporated in the model.<br>**Download URL:** https://zenodo.org/record/8222698/files/pbmc_gedi_model_donor1_celltype.rds?download=1 |
| **File name:** pbmc_gedi_model_donor2_celltype.rds (related to Figure 4)<br>**Description:** GEDI object of PBMC data (donor 2), with prior information of cell type signatures incorporated in the model.<br>**Download URL:** https://zenodo.org/record/8222698/files/pbmc_gedi_model_donor2_celltype.rds?download=1 |
| **File name:** pbmc_gedi_model_bothDonors_TF.rds (related to Figure 4)<br>**Description:** GEDI object of PBMC data, with prior information of transcription factor regulatory networks incorporated in the model.<br>**Download URL:** https://zenodo.org/record/8222698/files/pbmc_gedi_model_bothDonors_TF.rds?download=1 |
| **File name:** COVID19_gedi_model_cohort1_TF.rds (related to Figure 4)<br>**Description:** GEDI object of COVID-19 data (cohort1), with prior information of transcription factor regulatory networks and sample-level variables both incorporated in the model.<br>**Download URL:** https://zenodo.org/record/8222698/files/COVID19_gedi_model_cohort1_TF.rds?download=1 |
| **File name:** Tasic_gedi_model.rds (related to Figure 5)<br>**Description:** GEDI object of Tasic dataset.<br>**Download URL:** https://zenodo.org/record/8222698/files/Tasic_gedi_model.rds?download=1 |
| **File name:** Faure_gedi_model.rds (related to Figure 5)<br>**Description:** GEDI object of Faure dataset.<br>**Download URL:** https://zenodo.org/record/8222698/files/Faure_gedi_model.rds?download=1 |
| **File name:** LaManno_gedi_model.rds (related to Figure 5)<br>**Description:** GEDI object of La Manno data, with prior information of post-transcriptional regulatory networks incorporated in the model.<br>**Download URL:** https://zenodo.org/record/8222698/files/LaManno_gedi_model.rds?download=1 |
