## Supplementary Methods for "A unified model for interpretable latent embedding of multi-sample, multi-condition single-cell data"

### 1 The GEDI framework

#### 1.1 Fitting the GEDI model with no prior gene-level or sample-level information

We start by discussing the solution for the simplest GEDI model, i.e., a model in which measurement matrix $\boldsymbol{Y}\in\mathbb{R}^{G\times N}$ is directly provided, there is no prior gene-level information, and there is no prior sample-level information. In this case, we aim to obtain the maximum a posteriori (MAP) estimate of the parameter set Θ:

$$\Theta = \{\boldsymbol{o}_r, \boldsymbol{Z}_r, \Delta\boldsymbol{o}_1, \dots, \Delta\boldsymbol{o}_Q, \Delta\boldsymbol{Z}_1, \dots, \Delta\boldsymbol{Z}_Q, \boldsymbol{B}, \boldsymbol{s}, \sigma^2\}$$

$$\widehat{\Theta}_{\text{MAP}}(\boldsymbol{Y}) = \arg\max_{\Theta} f(\boldsymbol{Y}|\Theta)g(\Theta)$$

Here, $Q$ represents the total number of datasets that are being integrated, $f$ is the sampling distribution of $\boldsymbol{Y}$ conditional on Θ, and $g$ is the prior distribution of Θ. Other parameters are the same as defined in the main **Methods** text.
$\boldsymbol{Y}$ consists of $N$ column vectors $\boldsymbol{y}_{n\in\{1,\dots,N\}}$; each vector $\boldsymbol{y}_n$ represents the measurements of $G$ genes/events in cell $n$. We assume that each observation $\boldsymbol{y}_n$ is independent of other observations conditional on Θ. Therefore:

$$\boldsymbol{y}_n|\Theta \sim \mathcal{N}\left(\boldsymbol{o}_r + \Delta\boldsymbol{o}_{i(n)} + \left(\boldsymbol{Z}_r + \Delta\boldsymbol{Z}_{i(n)}\right)\boldsymbol{b}_n + s_n\boldsymbol{1}_G, \sigma^2\boldsymbol{I}\right) \tag{1}$$

The column vector $\boldsymbol{o}_r\in\mathbb{R}^G$ represents the origin point on a reference hyperplane, with each element $o_{r,g}$ ($g\in\{1,\dots,G\}$) derived from the following prior:

$$o_{r,g} \sim \mathcal{N}(\mu_o, \sigma^2 S_o\boldsymbol{I})$$

$\Delta\boldsymbol{o}_i$ represents the sample-specific translation of the origin point. These vectors have the following priors:

$$\Delta\boldsymbol{o}_i \sim \mathcal{N}\left(\boldsymbol{0}, \sigma^2 S_{\Delta o_i}\boldsymbol{I}\right)$$

Each column $\boldsymbol{z}_{r,k}$ of the matrix $\boldsymbol{Z}_r\in\mathbb{R}^{G\times K}$ represents a vector that originates from point $\boldsymbol{o}_r$ and lies on the reference hyperplane, and has the following prior:

$$\boldsymbol{z}_{r,k} \sim \mathcal{N}(\boldsymbol{0}, \sigma^2 S_Z\boldsymbol{I})$$

Each column $\Delta\boldsymbol{z}_{i,k}$ of each matrix $\Delta\boldsymbol{Z}_i\in\mathbb{R}^{G\times K}$ (for all $i\in\{1,\dots,Q\}$) represents the sample-specific distortion of $\boldsymbol{z}_{r,k}$, and has the following prior:

$$\Delta\boldsymbol{z}_{i,k} \sim \mathcal{N}\left(\boldsymbol{0}, \sigma^2 S_{\Delta Z_i}\boldsymbol{I}\right)$$

Finally, each $s_n$ ($n\in\{1,\dots,N\}$) is a cell-specific intercept (representing library size) with the following prior:

$$s_n \sim N\left(\mu_{s,n}, \sigma_s^2\right)$$

The above priors include hyperparameters $\mu_o$, $S_o$, $S_{\Delta oi}$, $S_Z$, $S_{\Delta Zi}$, $\mu_{s,n}$ and $\sigma_s^2$. The choice of these hyperparameters are discussed in **section 1.6**.

The MAP estimate of the parameter set can be obtained as follows:

$$\begin{aligned}
\widehat{\Theta}_{\text{MAP}}(\boldsymbol{Y}) &= \arg\max_{\Theta} f(\boldsymbol{Y}|\Theta)g(\Theta) = \arg\min_{\Theta}[-\log[f(\boldsymbol{Y}|\Theta)g(\Theta)]] \\
&= \arg\min_{\Theta}\Bigg[\frac{GN}{2}log(2\pi\sigma^2) + \frac{1}{2\sigma^2}\sum_{n\in\{1,\dots,N\}}\left\|\boldsymbol{y}_n - \boldsymbol{o}_r - \Delta\boldsymbol{o}_{i(n)} - \left(\boldsymbol{Z}_r + \Delta\boldsymbol{Z}_{i(n)}\right)\boldsymbol{b}_n - s_n\boldsymbol{1}_G\right\|_2^2 \\
&+ \frac{G}{2}log(2\pi\sigma^2 S_o) + \frac{1}{2\sigma^2 S_o}\|\boldsymbol{o}_r - \mu_o\boldsymbol{1}_G\|_2^2 + \sum_{i\in\{1,\dots,Q\}}\left[\frac{G}{2}log\left(2\pi\sigma^2 S_{\Delta o_i}\right) + \frac{1}{2\sigma^2 S_{\Delta o_i}}\|\Delta\boldsymbol{o}_i\|_2^2\right] \\
&+ \frac{GK}{2}log(2\pi\sigma^2 S_Z) + \frac{1}{2\sigma^2 S_Z}\|\boldsymbol{Z}_r\|_F^2 + \sum_{i\in\{1,\dots,Q\}}\left[\frac{GK}{2}log\left(2\pi\sigma^2 S_{\Delta z_i}\right) + \frac{1}{2\sigma^2 S_{\Delta z_i}}\|\Delta\boldsymbol{Z}_i\|_F^2\right] \\
&+ \frac{1}{2\sigma_s^2}\sum_{n\in\{1,\dots,N\}}\left(s_n - \mu_{s,n}\right)^2\Bigg]
\end{aligned} \tag{2}$$

We use block coordinate descent to iteratively optimize the parameters until convergence, as described below.

##### 1.1.1 Solving $\boldsymbol{Z}_r$

From Eq. (2), if we isolate the terms that contain $\boldsymbol{Z}_r$, we have:

$$\widehat{\boldsymbol{Z_r}} = \arg\min_{\boldsymbol{Z}_r}\left[\frac{1}{2\sigma^2}\sum_{n\in\{1,\dots,N\}}\left\|\boldsymbol{y}_n - \boldsymbol{o}_r - \Delta\boldsymbol{o}_{i(n)} - \left(\boldsymbol{Z}_r + \Delta\boldsymbol{Z}_{i(n)}\right)\boldsymbol{b}_n - s_n\boldsymbol{1}_G\right\|_2^2 + \frac{1}{2\sigma^2 S_Z}\|\boldsymbol{Z}_r\|_F^2\right]$$
$$= \arg\min_{\boldsymbol{Z}_r}\left[\sum_{n\in\{1,\dots,N\}}\left\|\boldsymbol{y}_n - \boldsymbol{o}_r - \Delta\boldsymbol{o}_{i(n)} - \left(\boldsymbol{Z}_r + \Delta\boldsymbol{Z}_{i(n)}\right)\boldsymbol{b}_n - s_n\boldsymbol{1}_G\right\|_2^2 + \frac{1}{S_Z}\|\boldsymbol{Z}_r\|_F^2\right]$$

When $\boldsymbol{Z}_r$ is not restricted to be orthogonal, we can simply solve it as:

$$\widehat{\boldsymbol{Z_r}} = \boldsymbol{Y}'\boldsymbol{B}^T\left(\boldsymbol{B}\boldsymbol{B}^T + \frac{1}{S_Z}\boldsymbol{I}\right)^{-1}$$

where

$$\boldsymbol{Y}' = [\boldsymbol{y}_1' \quad \dots \quad \boldsymbol{y}_N']$$
$$\boldsymbol{y}_n' = \boldsymbol{y}_n - \boldsymbol{o}_r - \Delta\boldsymbol{o}_{i(n)} - \Delta\boldsymbol{Z}_{i(n)}\boldsymbol{b}_n - s_n\boldsymbol{1}_G \tag{3}$$

When $\boldsymbol{Z}_r$ is restricted so that its columns are orthogonal to each other, we solve the problem by first defining $\boldsymbol{Z}_r$ as the product of two matrices $\boldsymbol{U}\in\mathbb{R}^{G\times K}$ and $\boldsymbol{S}\in\mathbb{R}^{K\times K}$, where $\boldsymbol{U}$ is a matrix with orthonormal columns and $\boldsymbol{S}$=diag($\boldsymbol{s}$) is a diagonal matrix. This leads to the following minimization problem:

$$\widehat{\boldsymbol{Z_r}} = \widehat{\boldsymbol{U}}\widehat{\boldsymbol{S}}$$
$$\left\{\widehat{\boldsymbol{U}}, \widehat{\boldsymbol{S}}\right\} = \arg\min_{\{\boldsymbol{U},\boldsymbol{S}\}}\left\{\|\boldsymbol{Y}' - \boldsymbol{USB}\|_F^2 + \frac{1}{S_Z}\|\boldsymbol{US}\|_F^2\right\} = \arg\min_{\{\boldsymbol{U},\boldsymbol{S}\}}\|\boldsymbol{Y}'' - \boldsymbol{USB}'\|_F^2$$

where:

$$\boldsymbol{Y}'' = [\boldsymbol{Y}' \quad \boldsymbol{0}_{G\times K}]$$
$$\boldsymbol{B}' = \left[\boldsymbol{B} \quad \frac{1}{\sqrt{S_Z}}\boldsymbol{I}_{K\times K}\right]$$

We solve $\boldsymbol{U}$ and $\boldsymbol{S}$ iteratively as part of the block coordinate descent algorithm. First, to solve $\boldsymbol{S}$, we want to obtain:

$$\widehat{\boldsymbol{S}} = \arg\min_{\boldsymbol{S}}\|\boldsymbol{Y}'' - \boldsymbol{USB}'\|_2^2 = \arg\min_{\boldsymbol{S}}\sum_{g=1}^{G}\sum_{n=1}^{N+K}\left(y_{g,n}'' - \sum_{k=1}^{K}u_{g,k}s_k b_{k,n}'\right)^2$$
$$= \arg\min_{\boldsymbol{S}}\sum_g\sum_n\left(y_{g,n}''\right)^2 - 2\sum_g\sum_n\left(y_{g,n}''\sum_k u_{g,k}s_k b_{k,n}'\right) + \sum_g\sum_n\left(\sum_k u_{g,k}s_k b_{k,n}'\right)^2$$

For each element $s_\kappa$, we can separately solve by setting its derivative to zero:

$$\frac{d}{d(s_\kappa)}\left[\sum_g\sum_n\left(y_{g,n}''\right)^2 - 2\sum_g\sum_n\left(y_{g,n}''\sum_k u_{g,k}s_k b_{k,n}'\right) + \sum_g\sum_n\left(\sum_k u_{g,k}s_k b_{k,n}'\right)^2\right] = 0$$
$$\Rightarrow -2\sum_g\sum_n\left(y_{g,n}''u_{g,\kappa}b_{\kappa,n}'\right) + 2\sum_g\sum_n u_{g,\kappa}b_{\kappa,n}'\sum_k u_{g,k}s_k b_{k,n}' = 0$$
$$\Rightarrow -\sum_g\sum_n\left(y_{g,n}''u_{g,\kappa}b_{\kappa,n}'\right) + s_\kappa\sum_g\sum_n\left(u_{g,\kappa}b_{\kappa,n}'\right)^2 + \sum_g\sum_n u_{g,\kappa}b_{\kappa,n}'\sum_{k\neq\kappa}u_{g,k}s_k b_{k,n}' = 0$$

which can be rearranged as:

$$s_\kappa\sum_g\sum_n\left(u_{g,\kappa}b_{\kappa,n}\right)^2 = \sum_g\sum_n\left(y_{g,n}''u_{j,\kappa}b_{\kappa,n}'\right) - \sum_g\sum_n u_{g,\kappa}b_{\kappa,n}'\sum_{k\neq\kappa}u_{g,k}s_k b_{k,n}'$$
$$\Rightarrow s_\kappa = \frac{\sum_g\sum_n\left(y_{g,n}''u_{g,\kappa}b_{\kappa,n}'\right) - \sum_g\sum_n u_{g,\kappa}b_{\kappa,n}'\sum_{k\neq\kappa}u_{g,k}s_k b_{k,n}'}{\sum_g\sum_n\left(u_{g,\kappa}b_{\kappa,n}'\right)^2}$$
$$= \frac{\sum_g\sum_n\left(y_{g,n}''u_{g,\kappa}b_{\kappa,n}'\right) - \sum_n\sum_{k\neq\kappa}b_{\kappa,n}'s_k b_{k,n}'\sum_g u_{g,\kappa}u_{g,k}}{\sum_g\sum_n\left(u_{g,\kappa}b_{\kappa,n}'\right)^2}$$

Then, considering the fact that the different columns of $\boldsymbol{U}$ are orthonormal, and therefore their dot products are zero, this equation simplifies to:

$$s_\kappa = \frac{\sum_g \sum_n \left(y''_{g,n} u_{g,\kappa} b'_{\kappa,n}\right)}{\sum_g \sum_n \left(u_{g,\kappa} b'_{\kappa,n}\right)^2} = \frac{\sum_g u_{g,\kappa} \sum_n b'_{\kappa,n} y''_{g,n}}{\sum_n \left(b'_{\kappa,n}\right)^2 \sum_g \left(u_{g,\kappa}\right)^2} = \frac{\sum_g u_{g,\kappa} \sum_n b'_{\kappa,n} y''_{g,n}}{\sum_n \left(b'_{\kappa,n}\right)^2} = \frac{\sum_g u_{g,\kappa} \sum_n b'_{\kappa,n} y''_{g,n}}{1 + \frac{1}{S_Z}}$$
$$= \frac{\sum_j u_{j,\kappa} [\boldsymbol{Y}'(\boldsymbol{B}')^T]_{j,\kappa}}{1 + \frac{1}{S_Z}} \tag{4}$$

To solve $\boldsymbol{U}$, we have:

$$\widehat{\boldsymbol{U}} = \arg\min_{\boldsymbol{S}} \|\boldsymbol{Y}'' - \boldsymbol{USB}'\|_2^2$$

When $\boldsymbol{S}$ is given, this problem transforms into a Procrustes problem. Therefore, we can solve $\boldsymbol{U}$ by performing SVD on $\boldsymbol{Y}''(\boldsymbol{SB}')^T$:

$$\mathrm{SVD}[\boldsymbol{Y}''(\boldsymbol{SB}')^T] = \boldsymbol{U}'\boldsymbol{S}'\boldsymbol{V}^{\boldsymbol{T}}$$
$$\boldsymbol{U} = \boldsymbol{U}'\boldsymbol{V}^{\boldsymbol{T}} \tag{5}$$

###### 1.1.2 Solving $\Delta \boldsymbol{Z}_i$

For each given $i \in \{1,\ldots,Q\}$, we can isolate from Eq. (2) the terms that contain $\Delta \boldsymbol{Z}_i$:

$$\widehat{\Delta \boldsymbol{Z}_\iota} = \arg\min_{\Delta \boldsymbol{Z}_i} \left[ \frac{1}{2\sigma^2} \sum_{n \in \mathcal{D}_i} \|\boldsymbol{y}_n - \boldsymbol{o}_r - \Delta \boldsymbol{o}_i - (\boldsymbol{Z}_r + \Delta \boldsymbol{Z}_i)\boldsymbol{b}_n - s_n \mathbf{1}_G\|_2^2 + \frac{1}{2\sigma^2 S_{\Delta z_i}} \|\Delta \boldsymbol{Z}_i\|_F^2 \right]$$

Here, $\mathcal{D}_i$ is the set of indices for all cells that belong to sample (dataset) $i$, i.e., $\mathcal{D}_i \in \{n | i(n) = i\}$, where $i(n)$ is the sample to which cell $n$ belongs. We can solve $\Delta \boldsymbol{Z}_i$ as:

$$\widehat{\Delta \boldsymbol{Z}_\iota} = \boldsymbol{Y}'_i \boldsymbol{B}_i^T \left( \boldsymbol{B}_i \boldsymbol{B}_i^T + \frac{1}{S_{\Delta z_i}} \boldsymbol{I} \right)^{-1} \tag{6}$$

Here, $\boldsymbol{B}_i$ is a submatrix containing only columns $\mathcal{D}_i$ of of $\boldsymbol{B}$ (i.e, the concatenation of $\boldsymbol{b}_n$ column vectors for all cells $n$ that belong to sample $i$). Similarly, $\boldsymbol{Y}'_i$ here is defined as the concatenation of column vectors $\boldsymbol{y}'_n$ for all cells $n$ that belong to sample $i$, with $\boldsymbol{y}'_n$ here defined as:

$$\boldsymbol{y}'_n = \boldsymbol{y}_n - \boldsymbol{o}_r - \Delta \boldsymbol{o}_{i(n)} - \boldsymbol{Z}_r \boldsymbol{b}_n - s_n \mathbf{1}_G$$

###### 1.1.3 Solving $\boldsymbol{o}_r$

Following the same procedure as the sections above, we have:

$$\widehat{\boldsymbol{o}_r} = \arg\min_{\boldsymbol{o}_r} \left[ \frac{1}{2\sigma^2} \sum_{n \in \{1,\ldots,N\}} \left\|\boldsymbol{y}_n - \boldsymbol{o}_r - \Delta \boldsymbol{o}_{i(n)} - \left(\boldsymbol{Z}_r + \Delta \boldsymbol{Z}_{i(n)}\right)\boldsymbol{b}_n - s_n \mathbf{1}_G\right\|_2^2 + \frac{1}{2\sigma^2 S_o} \|\boldsymbol{o}_r - \mu_o \mathbf{1}_G\|_2^2 \right]$$
$$\widehat{\boldsymbol{o}_r} = \frac{\boldsymbol{Y}'\mathbf{1}_N + \frac{1}{S_o}\mu_o \mathbf{1}_G}{N + \frac{1}{S_o}} \tag{7}$$

Here, $\boldsymbol{Y}'$ is defined as:

$$\boldsymbol{Y}' = [\boldsymbol{y}'_1 \quad \ldots \quad \boldsymbol{y}'_N]$$
$$\boldsymbol{y}'_n = \boldsymbol{y}_n - \Delta \boldsymbol{o}_{i(n)} - \left(\boldsymbol{Z}_r + \Delta \boldsymbol{Z}_{i(n)}\right)\boldsymbol{b}_n - s_n \mathbf{1}_G$$

###### 1.1.4 Solving $\Delta \boldsymbol{o}_i$

Similar to the previous sections, we can isolate the terms that contain $\Delta o$, which leads to the following minimization problem:

$$\widehat{\Delta \boldsymbol{o}_\iota} = \arg\min_{\Delta \boldsymbol{o}_i} \left[ \frac{1}{2\sigma^2} \sum_{n \in \mathcal{D}_i} \|\boldsymbol{y}_n - \boldsymbol{o}_r - \Delta \boldsymbol{o}_i - (\boldsymbol{Z}_r + \Delta \boldsymbol{Z}_i)\boldsymbol{b}_n - s_n \mathbf{1}_G\|_2^2 + \frac{1}{2\sigma^2 S_{\Delta o_i}} \|\Delta \boldsymbol{o}_i\|_2^2 \right]$$
$$\widehat{\boldsymbol{o}_r} = \frac{\boldsymbol{Y}'_i \mathbf{1}_{N_i}}{N_i + \frac{1}{S_{\Delta o_i}}} \tag{8}$$

Here, $N_i$ is the number of cells in sample $i$, i.e., $N_i=|\mathcal{D}_i|$, and $\boldsymbol{Y}'_i$ here is defined as the concatenatation of column vectors $\boldsymbol{y}'_n$ for all cells $n\in\mathcal{D}_i$, with $\boldsymbol{y}'_n$ here defined as:

$$\boldsymbol{y}'_n = \boldsymbol{y}_n - \boldsymbol{o}_r - (\boldsymbol{Z}_r + \Delta\boldsymbol{Z}_{i(n)})\boldsymbol{b}_n - s_n\mathbf{1}_G$$

##### 1.1.5 Solving $\boldsymbol{B}$

By isolating the terms containing $\boldsymbol{B}$ in Eq. (2), we have:

$$\widehat{\boldsymbol{B}} = \arg\min_{\boldsymbol{B}\in\mathcal{B}} \frac{1}{2\sigma^2} \sum_{n\in\{1,\ldots,N\}} \left\|\boldsymbol{y}_n - \boldsymbol{o}_r - \Delta\boldsymbol{o}_{i(n)} - (\boldsymbol{Z}_r + \Delta\boldsymbol{Z}_{i(n)})\boldsymbol{b}_n - s_n\mathbf{1}_G\right\|_2^2$$

Here, $\mathcal{B}\subseteq\mathbb{R}^{K\times N}$ is the subset of $\mathbb{R}^{K\times N}$ to which $\boldsymbol{B}$ is restricted. In its simplest form, we have:

$$\mathcal{B} = \left\{\mathbb{R}^{K\times N}\middle|\ \forall k \in \{1,\ldots,K\}\ \textstyle\sum_{n=1}^{N}\left(b_{k,n}\right)^2 = 1\right\}$$

We can solve separately for each $\boldsymbol{b}_n$:

$$\widehat{\boldsymbol{b}_n} = \arg\min_{\boldsymbol{b}_n} \left\|\boldsymbol{y}_n - \boldsymbol{o}_r - \Delta\boldsymbol{o}_{i(n)} - (\boldsymbol{Z}_r + \Delta\boldsymbol{Z}_{i(n)})\boldsymbol{b}_n - s_n\mathbf{1}_G\right\|_2^2$$

This leads us to:

$$\widehat{\boldsymbol{b}'_n} = \left(\boldsymbol{W}_{i(n)}^T\boldsymbol{W}_{i(n)}\right)^{-1}\boldsymbol{W}_{i(n)}^T\boldsymbol{y}'_n$$
$$\widehat{\boldsymbol{b}_n} = \widehat{\boldsymbol{b}'_n}\odot\boldsymbol{d} \tag{9}$$

Here, $\boldsymbol{W}_{i(n)}$ is defined as $\boldsymbol{W}_{i(n)}=\boldsymbol{Z}_r+\Delta\boldsymbol{Z}_{i(n)}$, and $\boldsymbol{y}'_n$ is defined as $\boldsymbol{y}'_n=\boldsymbol{y}_n-\boldsymbol{o}_r-\Delta\boldsymbol{o}_{i(n)}-s_n\mathbf{1}_G$. The operator $\odot$ is the Hadamard product, and $\boldsymbol{d}$ is a vector of normalization factors, so that each row of $\boldsymbol{B}$ becomes a unit vector.

With the added restriction that $\boldsymbol{B}$ should be restricted to the points on an ellipsoid, the above equation can be modified as follows:

$$\widehat{\boldsymbol{b}'_n} = \rho_n\left(\boldsymbol{W}_{i(n)}^T\boldsymbol{W}_{i(n)}\right)^{-1}\boldsymbol{W}_{i(n)}^T\boldsymbol{y}'_n$$
$$\widehat{\boldsymbol{b}_n} = \widehat{\boldsymbol{b}'_n}\odot\boldsymbol{d} \tag{10}$$

Here, $\rho_n$ is a cell-specific normalization factor so that each $\boldsymbol{b}'_n$ becomes a unit vector (and, therefore, all points $\boldsymbol{b}'_n$ lie on the surface of a unit ($K$–1)-sphere). As in above, $\boldsymbol{d}$ is a vector of normalization factors, so that each row of $\boldsymbol{B}$ becomes a unit vector, but it also represents the lengths of the semi-axes of the ellipsoid on which the solution for $\boldsymbol{B}$ lies.

##### 1.1.6 Solving $\sigma^2$

Let's start by defining the following sum:

$$\mathcal{S} = \sum_{n\in\{1,\ldots,N\}} \left\|\boldsymbol{y}_n - \boldsymbol{o}_r - \Delta\boldsymbol{o}_{i(n)} - (\boldsymbol{Z}_r + \Delta\boldsymbol{Z}_{i(n)})\boldsymbol{b}_n - s_n\mathbf{1}_G\right\|_2^2 + \frac{1}{S_o}\|\boldsymbol{o}_r - \mu_o\mathbf{1}_G\|_2^2 + \sum_{i\in\{1,\ldots,Q\}}\frac{1}{S_{\Delta o_i}}\|\Delta\boldsymbol{o}_i\|_2^2 + \frac{1}{S_Z}\|\boldsymbol{Z}_r\|_F^2 + \sum_{i\in\{1,\ldots,Q\}}\frac{1}{S_{\Delta z_i}}\|\Delta\boldsymbol{Z}_i\|_F^2$$

We can then rewrite Eq. (2) as:

$$\widehat{\sigma^2} = \arg\min_{\sigma^2}\left[\frac{GN}{2}log(2\pi\sigma^2) + \frac{G}{2}log(2\pi\sigma^2 S_o) + \frac{G}{2}\sum_{i\in\{1,\ldots,Q\}}log\left(2\pi\sigma^2 S_{\Delta o_i}\right) + \frac{GK}{2}log(2\pi\sigma^2 S_Z) + \frac{GK}{2}\sum_{i\in\{1,\ldots,Q\}}log\left(2\pi\sigma^2 S_{\Delta z_i}\right) + \frac{1}{2\sigma^2}\mathcal{S}\right]$$

We can solve $\sigma^2$ by setting its derivative to zero:

$$\frac{d}{d\sigma^2}\left[\frac{GN}{2}log(2\pi) + \frac{G}{2}log(2\pi\sigma^2 S_o) + \frac{G}{2}\sum_{i\in\{1,\ldots,Q\}}log\left(2\pi\sigma^2 S_{\Delta o_i}\right) + \frac{GK}{2}log(2\pi\sigma^2 S_Z) + \frac{GK}{2}\sum_{i\in\{1,\ldots,Q\}}log\left(2\pi\sigma^2 S_{\Delta z_i}\right) + \frac{1}{2\sigma^2}\mathcal{S}\right] = 0$$

$$\Rightarrow (GN+G+GQ+GK+GKQ)\frac{1}{\sigma^2}-\left(\frac{1}{\sigma^2}\right)^2\mathcal{S}=0$$
$$\Rightarrow \widehat{\sigma^2}=\frac{\mathcal{S}}{GN+G+GQ+GK+GKQ} \tag{11}$$

#### 1.2 Fitting the GEDI model with gene-level prior information

We can expresses $\boldsymbol{Z}_r$ as a probabilistic function of $\boldsymbol{C}\in\mathbb{R}^{G\times P}$, where $\boldsymbol{C}$ is a matrix representing gene-level prior information matrix. In this case, each column $\boldsymbol{z}_{r,k}$ of $\boldsymbol{Z}_r$ is treated as a latent variable with the following conditional probability function:

$$\boldsymbol{z}_{r,k}|\boldsymbol{a}_k\sim\mathcal{N}(\boldsymbol{C}\boldsymbol{a}_k,\sigma^2 S_Z\boldsymbol{I})$$
$$\boldsymbol{a}_k\sim\mathcal{N}(\boldsymbol{0},\sigma^2 S_A\boldsymbol{I})$$

The column vectors $\boldsymbol{a}_{k\in\{1,\dots,K\}}$ together form the matrix $\boldsymbol{A}\in\mathbb{R}^{P\times K}$. Therefore, the parameter set to be optimized will now include:

$$\Theta=\{\boldsymbol{o}_r,\boldsymbol{A},\Delta\boldsymbol{o}_1,\dots,\Delta\boldsymbol{o}_Q,\Delta\boldsymbol{Z}_1,\dots,\Delta\boldsymbol{Z}_Q,\boldsymbol{B},\sigma^2\}$$

We need to then optimize the following function:

$$\widehat{\Theta}_{\mathrm{MAP}}(\boldsymbol{Y})=\arg\max_\Theta f(\boldsymbol{Y}|\Theta)g(\Theta)=\arg\max_\Theta\int f(\boldsymbol{Y}|\Theta,\boldsymbol{Z}_r)f(\boldsymbol{Z}_r|\Theta)d\boldsymbol{Z}_r\,g(\Theta)$$

For simplicity, in what follows, we drop the subscript $r$ from $\boldsymbol{Z}_r$, and refer to it simply as $\boldsymbol{Z}$.

We can solve this optimization problem using expectation-maximization. First, at each iteration $t$, we define the expectation function as:

$$\begin{aligned}Q(\Theta|\Theta^{(t)})&=E_{\boldsymbol{Z}|\boldsymbol{Y},\Theta^{(t)}}[\log[f(\boldsymbol{Y}|\Theta,\boldsymbol{Z})f(\boldsymbol{Z}|\Theta)g(\Theta)]]=E_{\boldsymbol{Z}|\boldsymbol{Y},\Theta^{(t)}}[\log f(\boldsymbol{Y}|\Theta,\boldsymbol{Z})+\log f(\boldsymbol{Z}|\Theta)+\log\ g(\Theta)]\\&=E_{\boldsymbol{Z}|\boldsymbol{Y},\Theta^{(t)}}[\log f(\boldsymbol{Y}|\Theta,\boldsymbol{Z})+\log f(\boldsymbol{Z}|\Theta)]+\log\ g(\Theta)\\&=E_{\boldsymbol{Z}|\boldsymbol{Y},\Theta^{(t)}}[\log f(\boldsymbol{Y}|\Theta,\boldsymbol{Z})]+E_{\boldsymbol{Z}|\boldsymbol{Y},\Theta^{(t)}}[\log f(\boldsymbol{Z}|\Theta)]+\log\ g(\Theta)\end{aligned}$$

Then, we maximize the function $Q$ with respect to each parameter using block coordinate descent, as described below.

##### 1.2.1 Solving $\boldsymbol{A}$

We want to maximize $Q$ with respect to $\boldsymbol{A}$:

$$\widehat{\boldsymbol{A}}=\arg\max_{\boldsymbol{A}}Q(\Theta|\Theta^{(t)})=\arg\max_{\boldsymbol{A}}\left[E_{\boldsymbol{Z}|\boldsymbol{Y},\Theta^{(t)}}[\log f(\boldsymbol{Y}|\Theta,\boldsymbol{Z})]+E_{\boldsymbol{Z}|\boldsymbol{Y},\Theta^{(t)}}[\log f(\boldsymbol{Z}|\Theta)]+\log\ g(\Theta)\right]$$

Note that, conditional on $\boldsymbol{Z}$, $\boldsymbol{Y}$ is independent of $\boldsymbol{A}$. Therefore:

$$\begin{aligned}\widehat{\boldsymbol{A}}&=\arg\max_{\boldsymbol{A}}\left[E_{\boldsymbol{Z}|\boldsymbol{Y},\Theta^{(t)}}[\log f(\boldsymbol{Y}|\Theta,\boldsymbol{Z})]+E_{\boldsymbol{Z}|\boldsymbol{Y},\Theta^{(t)}}[\log f(\boldsymbol{Z}|\Theta)]+\log\ g(\Theta)\right]\\&=\arg\max_{\boldsymbol{A}}\left[E_{\boldsymbol{Z}|\boldsymbol{Y},\Theta^{(t)}}[\log f(\boldsymbol{Z}|\Theta)]+\log\ g(\Theta)\right]=\arg\min_{\boldsymbol{A}}\left[E_{\boldsymbol{Z}|\boldsymbol{Y},\Theta^{(t)}}[-\log f(\boldsymbol{Z}|\Theta)]-\log\ g(\Theta)\right]\\&=\arg\min_{\boldsymbol{A}}\left[E_{\boldsymbol{Z}|\boldsymbol{Y},\Theta^{(t)}}\left[\frac{1}{2\sigma^2 S_Z}\|\boldsymbol{Z}-\boldsymbol{C}\boldsymbol{A}\|_F^2\right]+\frac{1}{2\sigma^2 S_A}\|\boldsymbol{A}\|_F^2\right]\\&=\arg\min_{\boldsymbol{A}}\left[\frac{1}{S_Z}E_{\boldsymbol{Z}|\boldsymbol{Y},\Theta^{(t)}}\left[\sum_{g=1}^{G}\sum_{k=1}^{K}\left(z_{g,k}-(\boldsymbol{C}\boldsymbol{A})_{g,k}\right)^2\right]+\frac{1}{S_A}\|\boldsymbol{A}\|_F^2\right]\end{aligned}$$

This equation can be expanded as:

$$\widehat{\boldsymbol{A}}=\arg\min_{\boldsymbol{A}}\left[\frac{1}{S_Z}\sum_{g=1}^{G}\sum_{k=1}^{K}\left(E_{\boldsymbol{Z}|\boldsymbol{Y},\Theta^{(t)}}(z_{g,k}^2)-2(\boldsymbol{C}\boldsymbol{A})_{g,k}E_{\boldsymbol{Z}|\boldsymbol{Y},\Theta^{(t)}}(z_{g,k})+(\boldsymbol{C}\boldsymbol{A})_{g,k}^2\right)+\frac{1}{S_A}\|\boldsymbol{A}\|_F^2\right]$$

Since $E(X^2)=\mathrm{Var}(X)+[E(X)]^2$, we can rewrite the above equation as:

$$\widehat{\boldsymbol{A}} = \arg\min_{\boldsymbol{A}} \left[ \frac{1}{S_Z} \sum_{g=1}^{G} \sum_{k=1}^{K} \left[ Var_{\boldsymbol{Z}|\boldsymbol{Y},\Theta^{(t)}}\left(z_{g,k}\right) + \left[ E_{\boldsymbol{Z}|\boldsymbol{Y},\Theta^{(t)}}\left(z_{g,k}\right)\right]^2 - 2(\boldsymbol{CA})_{g,k} E_{\boldsymbol{Z}|\boldsymbol{Y},\Theta^{(t)}}\left(z_{g,k}\right) + (\boldsymbol{CA})_{g,k}^2 \right] + \frac{1}{S_A} \|\boldsymbol{A}\|_F^2 \right]$$

$$= \arg\min_{\boldsymbol{A}} \left[ \frac{1}{S_Z} \sum_{g=1}^{G} \sum_{k=1}^{K} Var_{\boldsymbol{Z}|\boldsymbol{Y},\Theta^{(t)}}\left(z_{g,k}\right) + \frac{1}{S_Z} \sum_{g=1}^{G} \sum_{k=1}^{K} \left( E_{\boldsymbol{Z}|\boldsymbol{Y},\Theta^{(t)}}\left(z_{g,k}\right) - (\boldsymbol{CA})_{g,k} \right)^2 + \frac{1}{S_A} \|\boldsymbol{A}\|_F^2 \right]$$

We note that the variance of each element of $\boldsymbol{Z}$, conditional on $\boldsymbol{Y}$ and the current estimates of the parameters $\Theta^{(t)}$, is independent of $\boldsymbol{A}$. Therefore:

$$\widehat{\boldsymbol{A}} = \arg\min_{\boldsymbol{A}} \left[ \frac{1}{S_Z} \sum_{g=1}^{G} \sum_{k=1}^{K} \left( E_{\boldsymbol{Z}|\boldsymbol{Y},\Theta^{(t)}}\left(z_{g,k}\right) - (\boldsymbol{CA})_{g,k} \right)^2 + \frac{1}{S_A} \|\boldsymbol{A}\|_F^2 \right] = \left( \boldsymbol{C}^T\boldsymbol{C} + \frac{S_Z}{S_A}\boldsymbol{I} \right)^{-1} \boldsymbol{C}^T E_{\boldsymbol{Z}|\boldsymbol{Y},\Theta^{(t)}}(\boldsymbol{Z}) \quad (12)$$

Solving this equation requires the calculation of the expectation (mean) of $\boldsymbol{Z}$ given $\boldsymbol{Y}$ and the current estimates of the parameters $\Theta^{(t)}$. When $\boldsymbol{Z}$ is not restricted by orthogonality constraint, this is simply a Bayesian linear regression problem, leading to the following expectation:

$$E_{\boldsymbol{Z}|\boldsymbol{Y},\Theta^{(t)}}(\boldsymbol{Z}) = \left( \boldsymbol{Y}'\left(\boldsymbol{B}^{(t)}\right)^T + \frac{1}{S_Z}\boldsymbol{CA}^{(t)} \right)\left( \boldsymbol{B}^{(t)}\left(\boldsymbol{B}^{(t)}\right)^T + \frac{1}{S_Z}\boldsymbol{I} \right)^{-1} \quad (13)$$

Here, $\boldsymbol{Y}' \in \mathbb{R}^{G\times N}$ consists of $N$ column vectors $\boldsymbol{y}'_n \in \mathbb{R}^G$, each defined as:

$$\boldsymbol{y}'_n = \boldsymbol{y}_n - \boldsymbol{o}_r^{(t)} - \Delta\boldsymbol{o}_{i(n)}^{(t)} - \Delta\boldsymbol{Z}_{i(n)}^{(t)}\boldsymbol{b}_n^{(t)} - s_n^{(t)}\boldsymbol{1}_G$$

Note that, since each column $\boldsymbol{z}_k$ of $\boldsymbol{Z}$ has a multivariate normal posterior distribution, its mean coincides with its mode. We use this property to approximate the mean of $\boldsymbol{Z}$ when it is restricted to have orthogonal columns; i.e., we replace $E_{\boldsymbol{Z}|\boldsymbol{Y},\Theta}(\boldsymbol{Z})$ with $\boldsymbol{Z}_{\mathrm{MAP}}(\boldsymbol{Y},\Theta^{(t)})$ with orthogonality constraint:

$$E_{\boldsymbol{Z}|\boldsymbol{Y},\Theta^{(t)}}(\boldsymbol{Z}) \cong \boldsymbol{Z}_{MAP}\left(\boldsymbol{Y},\Theta^{(t)}\right)$$

$$= \arg\min_{\boldsymbol{Z}|\forall k\neq k'\ (\boldsymbol{Z}^T\boldsymbol{Z})_{k,k'}=0} \left[ \frac{1}{2\sigma^2} \sum_{n\in\{1,\dots,N\}} \left\|\boldsymbol{y}_n - \boldsymbol{o}_r - \Delta\boldsymbol{o}_{i(n)} - \left(\boldsymbol{Z} + \Delta\boldsymbol{Z}_{i(n)}\right)\boldsymbol{b}_n - s_n\boldsymbol{1}_G\right\|_2^2 + \frac{1}{2\sigma^2 S_Z}\|\boldsymbol{Z} - \boldsymbol{CA}\|_F^2 \right]$$

$$= \arg\min_{\boldsymbol{Z}|\forall k\neq k'\ (\boldsymbol{Z}^T\boldsymbol{Z})_{k,k'}=0} \left[ \sum_{n\in\{1,\dots,N\}} \left\|\boldsymbol{y}_n - \boldsymbol{o}_r - \Delta\boldsymbol{o}_{i(n)} - \left(\boldsymbol{Z} + \Delta\boldsymbol{Z}_{i(n)}\right)\boldsymbol{b}_n - s_n\boldsymbol{1}_G\right\|_2^2 + \frac{1}{S_Z}\|\boldsymbol{Z} - \boldsymbol{CA}\|_F^2 \right]$$

This can be solved similar to **section 1.1.1**, with the only difference that, here, we define $\boldsymbol{Y}''$ as:

$$\boldsymbol{Y}'' = \left[ \boldsymbol{Y}' \quad \frac{1}{\sqrt{S_Z}}\boldsymbol{CA} \right]$$

##### 1.2.2 Solving $\sigma^2$

We maximize the function $Q$ with respect to $\sigma^2$:

$$\widehat{\sigma^2} = \arg\max_{\sigma^2} Q\left(\Theta\middle|\Theta^{(t)}\right) = \arg\max_{\sigma^2} \left[ E_{\boldsymbol{Z}|\boldsymbol{Y},\Theta^{(t)}}[\log f(\boldsymbol{Y}|\Theta,\boldsymbol{Z})] + E_{\boldsymbol{Z}|\boldsymbol{Y},\Theta^{(t)}}[\log f(\boldsymbol{Z}|\Theta)] + \log\ g(\Theta) \right]$$

$$= \arg\min_{\sigma^2} \left[ -E_{\boldsymbol{Z}|\boldsymbol{Y},\Theta^{(t)}}[\log f(\boldsymbol{Y}|\Theta,\boldsymbol{Z})] - E_{\boldsymbol{Z}|\boldsymbol{Y},\Theta^{(t)}}[\log f(\boldsymbol{Z}|\Theta)] - \log\ g(\Theta) \right]$$

Using a procedure similar to the previous section, we can show that:

$$-E_{\boldsymbol{Z}|\boldsymbol{Y},\Theta^{(t)}}[\log f(\boldsymbol{Z}|\Theta)] = \frac{GK}{2} log(2\pi\sigma^2 S_Z) + \frac{1}{2\sigma^2 S_Z} E_{\boldsymbol{Z}|\boldsymbol{Y},\Theta^{(t)}}\left[\sum_{g=1}^{G}\sum_{k=1}^{K}\left(z_{g,k} - (\boldsymbol{CA})_{g,k}\right)^2\right]$$
$$= \frac{GK}{2} log(2\pi\sigma^2 S_Z) + \frac{1}{2\sigma^2 S_Z}\sum_{g=1}^{G}\sum_{k=1}^{K}\left(E_{\boldsymbol{Z}|\boldsymbol{Y},\Theta^{(t)}}(z_{g,k}) - (\boldsymbol{CA})_{g,k}\right)^2$$
$$+ \frac{1}{2\sigma^2 S_Z}\sum_{g=1}^{G}\sum_{k=1}^{K} Var_{\boldsymbol{Z}|\boldsymbol{Y},\Theta^{(t)}}(z_{g,k})$$

The variance of each element of $\boldsymbol{Z}$, conditional on $\boldsymbol{Y}$ and the current estimates of the parameters $\Theta^{(t)}$, is given by:

$$Var_{\boldsymbol{Z}|\boldsymbol{Y},\Theta^{(t)}}(z_{g,k}) = \left(\sigma^{(t)}\right)^2\left[\left(\boldsymbol{B}^{(t)}\left(\boldsymbol{B}^{(t)}\right)^T + \frac{1}{S_Z}\boldsymbol{I}\right)^{-1}\right]_{k,k}$$

Therefore:

$$-E_{\boldsymbol{Z}|\boldsymbol{Y},\Theta^{(t)}}[\log f(\boldsymbol{Z}|\Theta)]$$
$$= \frac{GK}{2} log(2\pi\sigma^2 S_Z) + \frac{1}{2\sigma^2 S_Z}\sum_{g=1}^{G}\sum_{k=1}^{K}\left(E_{\boldsymbol{Z}|\boldsymbol{Y},\Theta^{(t)}}(z_{g,k}) - (\boldsymbol{CA})_{g,k}\right)^2$$
$$+ \frac{G\left(\sigma^{(t)}\right)^2}{2\sigma^2 S_Z}\sum_{k=1}^{K}\left[\left(\boldsymbol{B}^{(t)}\left(\boldsymbol{B}^{(t)}\right)^T + \frac{1}{S_Z}\boldsymbol{I}\right)^{-1}\right]_{k,k} \tag{14}$$

We can also use a similar approach to show that:

$$-E_{\boldsymbol{Z}|\boldsymbol{Y},\Theta^{(t)}}[\log f(\boldsymbol{Y}|\Theta,\boldsymbol{Z})] = \frac{GN}{2} log(2\pi\sigma^2) + \frac{1}{2\sigma^2} E_{\boldsymbol{Z}|\boldsymbol{Y},\Theta^{(t)}}\left[\sum_{g=1}^{G}\sum_{n=1}^{N}\left(y'_{g,n} - (\boldsymbol{Z}\boldsymbol{b}_n)_g\right)^2\right]$$
$$= \frac{GN}{2} log(2\pi\sigma^2) + \frac{1}{2\sigma^2}\sum_{n=1}^{N}\sum_{g=1}^{G}\left[y'_{g,n} - \sum_{k=1}^{K} b_{k,n} E_{\boldsymbol{Z}|\boldsymbol{Y},\Theta^{(t)}}(z_{g,k})\right]^2$$
$$+ \frac{1}{2\sigma^2}\sum_{n=1}^{N}\sum_{g=1}^{G} Var_{\boldsymbol{Z}|\boldsymbol{Y},\Theta^{(t)}}\left(\sum_{k=1}^{K} z_{g,k} b_{k,n}\right)$$

Here, $\boldsymbol{y}'_n$ is defined in the same way as in **section 1.2.1**. For the conditional variance of $\boldsymbol{Z}$, we have:

$$Var_{\boldsymbol{Z}|\boldsymbol{Y},\Theta^{(t)}}\left(\sum_{k=1}^{K} z_{g,k} b_{k,n}\right) = \sum_{k,k'=1}^{K} Cov_{\boldsymbol{Z}|\boldsymbol{Y},\Theta^{(t)}}(z_{g,k} b_{k,n}, z_{g,k'} b_{k',n}) = \sum_{k,k'=1}^{K} b_{k,n} b_{k',n} Cov_{\boldsymbol{Z}|\boldsymbol{Y},\Theta^{(t)}}(z_{g,k}, z_{g,k'})$$
$$= \left(\sigma^{(t)}\right)^2 \sum_{k,k'=1}^{K} b_{k,n} b_{k',n}\left[\left(\boldsymbol{B}^{(t)}\left(\boldsymbol{B}^{(t)}\right)^T + \frac{1}{S_Z}\boldsymbol{I}\right)^{-1}\right]_{k,k'}$$

Therefore:

$$-E_{\boldsymbol{Z}|\boldsymbol{Y},\Theta^{(t)}}[\log f(\boldsymbol{Y}|\Theta,\boldsymbol{Z})]$$
$$= \frac{GN}{2} log(2\pi\sigma^2) + \frac{1}{2\sigma^2}\sum_{n=1}^{N}\sum_{g=1}^{G}\left[y'_{g,n} - \sum_{k=1}^{K} b_{k,n} E_{\boldsymbol{Z}|\boldsymbol{Y},\Theta^{(t)}}(z_{g,k})\right]^2$$
$$+ \frac{G\left(\sigma^{(t)}\right)^2}{2\sigma^2}\sum_{n=1}^{N}\sum_{k,k'=1}^{K} b_{k,n} b_{k',n}\left[\left(\boldsymbol{B}^{(t)}\left(\boldsymbol{B}^{(t)}\right)^T + \frac{1}{S_Z}\boldsymbol{I}\right)^{-1}\right]_{k,k'} \tag{15}$$

Finally, for the prior function $g$, we have:

$$-\log\ g(\Theta) = \frac{G}{2}log(2\pi\sigma^2 S_o) + \frac{1}{2\sigma^2 S_o}\|\boldsymbol{o}_r - \mu_o \mathbf{1}_G\|_2^2 + \frac{G}{2}\sum_{i\in\{1,\dots,Q\}} log\left(2\pi\sigma^2 S_{\Delta o_i}\right) + \frac{1}{2\sigma^2}\sum_{i\in\{1,\dots,Q\}}\frac{1}{S_{\Delta o_i}}\|\Delta\boldsymbol{o}_i\|_2^2$$
$$+\frac{PK}{2}log(2\pi\sigma^2 S_A) + \frac{1}{2\sigma^2 S_A}\|\boldsymbol{A}\|_F^2 + \frac{GK}{2}\sum_{i\in\{1,\dots,Q\}} log\left(2\pi\sigma^2 S_{\Delta z_i}\right) + \frac{1}{2\sigma^2}\sum_{i\in\{1,\dots,Q\}}\frac{1}{S_{\Delta z_i}}\|\Delta\boldsymbol{Z}_i\|_F^2 \tag{16}$$

Now, let's define the following sum:

$$\mathcal{S} = \sum_{n=1}^{N}\sum_{g=1}^{G}\left[y'_{g,n} - \sum_{k=1}^{K} b_{k,n} E_{\boldsymbol{Z}|\boldsymbol{Y},\Theta^{(t)}}\left(z_{g,k}\right)\right]^2 + G\left(\sigma^{(t)}\right)^2\sum_{n=1}^{N}\sum_{k,k'=1}^{K} b_{k,n}b_{k',n}\left[\left(\boldsymbol{B}^{(t)}\left(\boldsymbol{B}^{(t)}\right)^T + \frac{1}{S_Z}\boldsymbol{I}\right)^{-1}\right]_{k,k'}$$
$$+\frac{1}{S_Z}\sum_{g=1}^{G}\sum_{k=1}^{K}\left(E_{\boldsymbol{Z}|\boldsymbol{Y},\Theta^{(t)}}\left(z_{g,k}\right) - (\boldsymbol{CA})_{g,k}\right)^2 + \frac{1}{S_Z}G\left(\sigma^{(t)}\right)^2\sum_{k=1}^{K}\left[\left(\boldsymbol{B}^{(t)}\left(\boldsymbol{B}^{(t)}\right)^T + \frac{1}{S_Z}\boldsymbol{I}\right)^{-1}\right]_{k,k}$$
$$+\frac{1}{S_o}\|\boldsymbol{o}_r - \mu_o\mathbf{1}_G\|_2^2 + \sum_{i\in\{1,\dots,Q\}}\frac{1}{S_{\Delta o_i}}\|\Delta\boldsymbol{o}_i\|_2^2 + \frac{1}{S_A}\|\boldsymbol{A}\|_F^2 + \sum_{i\in\{1,\dots,Q\}}\frac{1}{S_{\Delta z_i}}\|\Delta\boldsymbol{Z}_i\|_F^2 \tag{17}$$

Combining Eq. (14)-(17), we have:

$$\widehat{\sigma^2} = \frac{\mathcal{S}}{GN + PK + G + GQ + GK + GKQ} \tag{18}$$

##### 1.2.3 Solving other parameters

As Eq. (14)-(17) in **section 1.2.2** suggest, maximization of the expectation function with respect to all parameters other than $\boldsymbol{A}$ and $\sigma^2$ can be done in the same way as in **sections 1.1.2-1.1.5**, with the exception that $\boldsymbol{Z}_r$ is replaced with its expectation, $E_{\boldsymbol{Z}|\boldsymbol{Y},\Theta}(\boldsymbol{Z})$.

#### 1.3 Fitting the GEDI model with sample-level prior information

The sample-specific parameters that specify the distortions of the manifold in sample $i$, i.e, $\Delta\boldsymbol{o}_i$,$\Delta\boldsymbol{Z}_i$, can be expressed as probabilistic functions of $\boldsymbol{h}_i\in\mathbb{R}^L$, where $\boldsymbol{h}_i$ is a column vector whose elements represent the values of $L$ variables for sample $i$. In this case, $\Delta\boldsymbol{o}_i$ as well as each column $\Delta\boldsymbol{z}_{i,k}$ are treated as latent variables with the following conditional probability function:

$$\Delta\boldsymbol{o}_i|\boldsymbol{R}_o\sim\mathcal{N}\left(\boldsymbol{R}_o\boldsymbol{h}_i, \sigma^2 S_{\Delta o_i}\boldsymbol{I}\right)$$
$$\boldsymbol{R}_o\sim\mathcal{N}\left(\boldsymbol{0}, \sigma^2 S_{R_o}\boldsymbol{I}\right)$$
$$\Delta\boldsymbol{z}_{i,k}|\boldsymbol{R}_k\sim\mathcal{N}\left(\boldsymbol{R}_k\boldsymbol{h}_i, \sigma^2 S_{\Delta Z_i}\boldsymbol{I}\right)$$
$$\boldsymbol{R}_k\sim\mathcal{N}\left(\boldsymbol{0}, \sigma^2 S_{R_k}\boldsymbol{I}\right)$$

$\boldsymbol{R}_o\in\mathbb{R}^{G\times L}$ and $\boldsymbol{R}_k\in\mathbb{R}^{G\times L}$ are matrices that represent the effects of the $L$ variables on $\Delta\boldsymbol{o}_i$ and $\Delta\boldsymbol{z}_{i,k}$, respectively. Here, we will discuss how to solve this model in the absence of gene-level prior information, but the solutions from this section and **section 1.2** can be combined to solve the model in the presence of both gene-level and sample-level prior information. Here, the parameter set to be optimized includes:

$$\Theta = \{\boldsymbol{o}_r, \boldsymbol{Z}_r, \boldsymbol{R}_o, \boldsymbol{R}_1, \dots, \boldsymbol{R}_K, \boldsymbol{B}, \sigma^2\}$$

For simplicity, let's define a set of matrices $\boldsymbol{\Delta}_i$, where for each $i\in\{1,\dots,Q\}$, the matrix $\boldsymbol{\Delta}_i\in\mathbb{R}^{G\times(K+1)}$ is the concatenation of $\Delta\boldsymbol{Z}_i$ and $\Delta\boldsymbol{o}_i$: $\boldsymbol{\Delta}_i$=[$\Delta\boldsymbol{Z}_i$ $\Delta\boldsymbol{o}_i$]. Also, let's define matrix $\boldsymbol{\Delta}\in\mathbb{R}^{G\times Q(K+1)}$ to be the concatenation of all submatrices $\boldsymbol{\Delta}_i$, $\boldsymbol{\Delta}$=[$\boldsymbol{\Delta}_1$ … $\boldsymbol{\Delta}_Q$]. Therefore, we need to optimize the following function:

$$\widehat{\Theta}_{\mathrm{MAP}}(\boldsymbol{Y}) = \arg\max_\Theta f(\boldsymbol{Y}|\Theta)g(\Theta) = \arg\max_\Theta \int f(\boldsymbol{Y}|\Theta,\boldsymbol{\Delta})f(\boldsymbol{\Delta}|\Theta)d\boldsymbol{\Delta}\, g(\Theta)$$

Similar to **section 2**, we will use expectation maximization, where the expectation function $Q$ is defined as:

$$Q\left(\Theta\middle|\Theta^{(t)}\right) = E_{\boldsymbol{\Delta}|\boldsymbol{Y},\Theta^{(t)}}[\log f(\boldsymbol{Y}|\Theta,\boldsymbol{\Delta})] + E_{\boldsymbol{\Delta}|\boldsymbol{Y},\Theta^{(t)}}[\log f(\boldsymbol{\Delta}|\Theta)] + \log\ g(\Theta)$$

For simplicity, let's also define matrices $\boldsymbol{B}'$ and $\boldsymbol{B}'_i$ as:

$$\boldsymbol{B}' = \begin{bmatrix} \boldsymbol{B} \\ \boldsymbol{1}_{\boldsymbol{N}}^T \end{bmatrix} \quad \boldsymbol{B}'_i = \begin{bmatrix} \boldsymbol{B}_i \\ \boldsymbol{1}_{N_i}^T \end{bmatrix} \quad i \in \{1, \dots, Q\}$$

Here, $\mathbf{1}_N$ and $\mathbf{1}_{N_i}$ are $1\times N$ and $1\times N_i$ column-vectors of 1s, respectively, where $N_i$ is the size of dataset $i$. We can now see that:

$$-E_{\boldsymbol{\Delta}|\boldsymbol{Y},\Theta^{(t)}}[\log f(\boldsymbol{Y}|\Theta,\boldsymbol{\Delta})] = \frac{GN}{2} log(2\pi\sigma^2) + \frac{1}{2\sigma^2} E_{\boldsymbol{\Delta}|\boldsymbol{Y},\Theta^{(t)}}\left[\sum_{g=1}^{G}\sum_{n=1}^{N}\left(y'_{g,n} - (\boldsymbol{\Delta b}'_n)_g\right)^2\right]$$

Here, $\boldsymbol{y'}_n$ is defined as:

$$\boldsymbol{y}'_n = \boldsymbol{y}_n - \boldsymbol{o}_r^{(t)} - \boldsymbol{Z}_r^{(t)}\boldsymbol{b}_n^{(t)} - s_n^{(t)}\mathbf{1}_G$$

Similar to section 1.2.2, we can show that:

$$\begin{aligned} &-E_{\boldsymbol{\Delta}|\boldsymbol{Y},\Theta^{(t)}}[\log f(\boldsymbol{Y}|\Theta,\boldsymbol{\Delta})] \\ &= \frac{GN}{2} log(2\pi\sigma^2) + \frac{1}{2\sigma^2}\sum_{n=1}^{N}\sum_{g=1}^{G}\left[y'_{g,n} - \sum_{k=1}^{K+1} b'_{k,n} E_{\boldsymbol{\Delta}|\boldsymbol{Y},\Theta^{(t)}}\left(\delta_{i(n),g,k}\right)\right]^2 \\ &+ \frac{G\left(\sigma^{(t)}\right)^2}{2\sigma^2}\sum_{n=1}^{N}\sum_{k,k'=1}^{K} b'_{k,n}b'_{k',n}\left[\left({\boldsymbol{B}'_{i(n)}}^{(t)}\left({\boldsymbol{B}'_{i(n)}}^{(t)}\right)^T + \boldsymbol{\Lambda}_{i(n)}\right)^{-1}\right]_{k,k'} \end{aligned} \tag{19}$$

Here, the prior precision matrix $\boldsymbol{\Lambda}_i$ is defined as:

$$\boldsymbol{\Lambda}_i = \begin{bmatrix} \frac{1}{S_{\Delta z_i}}\boldsymbol{I}_{K\times K} & \mathbf{0}_K \\ \mathbf{0}_K^T & \frac{1}{S_{\Delta o_i}} \end{bmatrix}$$

Also, similar to section 1.2.2, we can write:

$$\begin{aligned} &-E_{\boldsymbol{\Delta}|\boldsymbol{Y},\Theta^{(t)}}[\log f(\boldsymbol{\Delta}|\Theta)] \\ &= \frac{GK}{2}\sum_{i=1}^{Q}\log\left(2\pi\sigma^2 S_{\Delta z_i}\right) + \sum_{i=1}^{Q}\frac{1}{2\sigma^2 S_{\Delta z_i}} E_{\boldsymbol{\Delta}|\boldsymbol{Y},\Theta^{(t)}}\left[\sum_{g=1}^{G}\sum_{k=1}^{K}\left(\Delta z_{i,g,k} - (\boldsymbol{R}_k\boldsymbol{h}_i)_g\right)^2\right] \\ &+ \frac{G}{2}\sum_{i=1}^{Q}\log\left(2\pi\sigma^2 S_{\Delta o_i}\right) + \sum_{i=1}^{Q}\frac{1}{2\sigma^2 S_{\Delta o_i}} E_{\boldsymbol{\Delta}|\boldsymbol{Y},\Theta^{(t)}}\left[\sum_{g=1}^{G}\left(\Delta o_{i,g} - (\boldsymbol{R}_o\boldsymbol{h}_i)_g\right)^2\right] \end{aligned}$$

This can be further expanded to:

$$\begin{aligned} &-E_{\boldsymbol{\Delta}|\boldsymbol{Y},\Theta^{(t)}}[\log f(\boldsymbol{\Delta}|\Theta)] \\ &= \frac{GK}{2}\sum_{i=1}^{Q}\log\left(2\pi\sigma^2 S_{\Delta z_i}\right) + \sum_{i=1}^{Q}\frac{1}{2\sigma^2 S_{\Delta z_i}}\sum_{g=1}^{G}\sum_{k=1}^{K}\left(E_{\boldsymbol{\Delta}|\boldsymbol{Y},\Theta^{(t)}}\left(\Delta z_{i,g,k}\right) - (\boldsymbol{R}_k\boldsymbol{h}_i)_g\right)^2 \\ &+ \frac{G}{2}\sum_{i=1}^{Q}\log\left(2\pi\sigma^2 S_{\Delta o_i}\right) + \sum_{i=1}^{Q}\frac{1}{2\sigma^2 S_{\Delta o_i}}\sum_{g=1}^{G}\left(E_{\boldsymbol{\Delta}|\boldsymbol{Y},\Theta^{(t)}}\left(\Delta o_{i,g}\right) - (\boldsymbol{R}_o\boldsymbol{h}_i)_g\right)^2 \\ &+ \sum_{i=1}^{Q}\frac{1}{2\sigma^2 S_{\Delta z_i}}\sum_{g=1}^{G}\sum_{k=1}^{K} Var_{\boldsymbol{\Delta}|\boldsymbol{Y},\Theta^{(t)}}\left(\Delta z_{i,g,k}\right) + \sum_{i=1}^{Q}\frac{1}{2\sigma^2 S_{\Delta o_i}}\sum_{g=1}^{G} Var_{\boldsymbol{\Delta}|\boldsymbol{Y},\Theta^{(t)}}\left(\Delta o_{i,g}\right) \end{aligned}$$

where:

$$Var_{\boldsymbol{\Delta}|\boldsymbol{Y},\Theta^{(t)}}\left(\Delta z_{i,g,k}\right) = \left(\sigma^{(t)}\right)^2\left[\left(\boldsymbol{B}_i^{(t)}\left(\boldsymbol{B}_i^{(t)}\right)^T + \frac{1}{S_{\Delta z_i}}\boldsymbol{I}\right)^{-1}\right]_{k,k}$$

$$Var_{\boldsymbol{\Delta}|\boldsymbol{Y},\Theta^{(t)}}\left(\Delta o_{i,g}\right) = \frac{\left(\sigma^{(t)}\right)^2}{N_i + \frac{1}{S_{\Delta o_i}}}$$

Therefore:

$$
\begin{aligned}
-E_{\boldsymbol{\Delta}|\boldsymbol{Y},\Theta^{(t)}}[\log f(\boldsymbol{\Delta}|\Theta)]
&= \frac{GK}{2}\sum_{i=1}^{Q}\log\left(2\pi\sigma^2 S_{\Delta Z_i}\right) + \frac{1}{2\sigma^2}\sum_{i=1}^{Q}\frac{1}{S_{\Delta Z_i}}\sum_{g=1}^{G}\sum_{k=1}^{K}\left(E_{\boldsymbol{\Delta}|\boldsymbol{Y},\Theta^{(t)}}\left(\Delta z_{i,g,k}\right) - (\boldsymbol{R}_k\boldsymbol{h}_i)_g\right)^2 \\
&+ \frac{G}{2}\sum_{i=1}^{Q}\log\left(2\pi\sigma^2 S_{\Delta o_i}\right) + \frac{1}{2\sigma^2}\sum_{i=1}^{Q}\frac{1}{S_{\Delta o_i}}\sum_{g=1}^{G}\left(E_{\boldsymbol{\Delta}|\boldsymbol{Y},\Theta^{(t)}}\left(\Delta o_{i,g}\right) - (\boldsymbol{R}_o\boldsymbol{h}_i)_g\right)^2 \\
&+ \frac{G\left(\sigma^{(t)}\right)^2}{2\sigma^2}\sum_{i=1}^{Q}\frac{1}{S_{\Delta Z_i}}\sum_{k=1}^{K}\left[\left(\boldsymbol{B}_i^{(t)}\left(\boldsymbol{B}_i^{(t)}\right)^T + \frac{1}{S_{\Delta Z_i}}\boldsymbol{I}\right)^{-1}\right]_{k,k} + \frac{G\left(\sigma^{(t)}\right)^2}{2\sigma^2}\sum_{i=1}^{Q}\frac{1}{S_{\Delta o_i}\left(N_i + \frac{1}{S_{\Delta o_i}}\right)}
\end{aligned}
\tag{20}
$$

Finally, for the prior function $g$, we have:

$$
\begin{aligned}
-\log\, g(\Theta) &= \frac{G}{2}log(2\pi\sigma^2 S_o) + \frac{1}{2\sigma^2 S_o}\|\boldsymbol{o}_r - \mu_o\mathbf{1}_G\|_2^2 + \frac{GL}{2}log\left(2\pi\sigma^2 S_{R_o}\right) + \frac{1}{2\sigma^2 S_{R_o}}\|\boldsymbol{R}_o\|_2^2 + \frac{GK}{2}log(2\pi\sigma^2 S_Z) \\
&+ \frac{1}{2\sigma^2 S_Z}\|\boldsymbol{Z}_r\|_F^2 + \frac{GL}{2}\sum_{k\in\{1,\ldots,K\}} log\left(2\pi\sigma^2 S_{R_k}\right) + \frac{1}{2\sigma^2}\sum_{k\in\{1,\ldots,K\}}\frac{1}{S_{R_k}}\|\boldsymbol{R}_k\|_F^2
\end{aligned}
\tag{21}
$$

Note that the equations above require calculation of the expectation for $\boldsymbol{\Delta}_i=[\Delta\boldsymbol{Z}_i\ \Delta\boldsymbol{o}_i]$, which is given by the equation below:

$$
E_{\boldsymbol{\Delta}|\boldsymbol{Y},\Theta^{(t)}}(\boldsymbol{\Delta}_i) = \left(\boldsymbol{Y}'\left(\boldsymbol{B}'^{(t)}_i\right)^T + \frac{1}{S_{\Delta Z_i}}\boldsymbol{R}_k^{(t)}\boldsymbol{h}_i\right)\left(\boldsymbol{B}'^{(t)}_i\left(\boldsymbol{B}'^{(t)}_i\right)^T + \frac{1}{S_{\Delta Z_i}}\boldsymbol{I}\right)^{-1}
\tag{22}
$$

##### 1.3.1 Solving $\boldsymbol{R_o}$

Let's first define matrices $\boldsymbol{H'}$ and $\Delta\boldsymbol{O'}$ as follows:

$$
\boldsymbol{H}'_{\Delta\boldsymbol{O}} = \left[\frac{1}{\sqrt{S_{\Delta o_1}}}\boldsymbol{h}_1 \quad \cdots \quad \frac{1}{\sqrt{S_{\Delta o_Q}}}\boldsymbol{h}_Q\right]
$$

$$
\Delta\boldsymbol{O}' = \left[\frac{1}{\sqrt{S_{\Delta o_1}}}E_{\boldsymbol{\Delta}|\boldsymbol{Y},\Theta^{(t)}}(\Delta\boldsymbol{o}_1) \quad \cdots \quad \frac{1}{\sqrt{S_{\Delta o_Q}}}E_{\boldsymbol{\Delta}|\boldsymbol{Y},\Theta^{(t)}}\left(\Delta\boldsymbol{o}_Q\right)\right]
$$

Then, combining Eq. (19)-(21) and minimizing with respect to $\boldsymbol{R}_o$, we can see that:

$$
\boldsymbol{R}_o = \Delta\boldsymbol{O}'\boldsymbol{H}'^{\,T}_{\Delta\boldsymbol{O}}\left(\boldsymbol{H}'_{\Delta\boldsymbol{O}}\boldsymbol{H}'^{\,T}_{\Delta\boldsymbol{O}} + \frac{1}{S_{R_o}}\boldsymbol{I}\right)^{-1}
\tag{23}
$$

##### 1.3.2 Solving $\boldsymbol{R_k}$

Let's first redefine matrix $\boldsymbol{H'}$ as follows:

$$
\boldsymbol{H}'_{\Delta\boldsymbol{Z}} = \left[\frac{1}{\sqrt{S_{\Delta Z_1}}}\boldsymbol{h}_1 \quad \cdots \quad \frac{1}{\sqrt{S_{\Delta Z_Q}}}\boldsymbol{h}_Q\right]
$$

Also, for each $k\in\{1,\ldots,Q\}$, we will define the matrix $\Delta\boldsymbol{Z}'_k$ as follows:

$$
\Delta\boldsymbol{Z}'_k = \left[\frac{1}{\sqrt{S_{\Delta Z_1}}}E_{\boldsymbol{\Delta}|\boldsymbol{Y},\Theta^{(t)}}\left(\Delta\boldsymbol{z}_{1,k}\right) \quad \cdots \quad \frac{1}{\sqrt{S_{\Delta Z_Q}}}E_{\boldsymbol{\Delta}|\boldsymbol{Y},\Theta^{(t)}}\left(\Delta\boldsymbol{z}_{Q,k}\right)\right]
$$

Combining Eq. (19)-(21) and minimizing with respect to each $\boldsymbol{R}_k$, we can see that:

$$
\boldsymbol{R}_k = \Delta\boldsymbol{Z}'_k\boldsymbol{H}'^{\,T}_{\Delta\boldsymbol{Z}}\left(\boldsymbol{H}'_{\Delta\boldsymbol{Z}}\boldsymbol{H}'^{\,T}_{\Delta\boldsymbol{Z}} + \frac{1}{S_{R_k}}\boldsymbol{I}\right)^{-1}
\tag{24}
$$

###### 1.3.3 Solving $\sigma^2$

Let's define the following sum:

$$\mathcal{S} = \sum_{n=1}^{N}\sum_{g=1}^{G}\left[y'_{g,n} - \sum_{k=1}^{K+1} b'_{k,n} E_{\boldsymbol{\Delta}|\boldsymbol{Y},\Theta^{(t)}}\left(\delta_{i(n),g,k}\right)\right]^2$$
$$+ G\left(\sigma^{(t)}\right)^2 \sum_{n=1}^{N}\sum_{k,k'=1}^{K} b'_{k,n}b'_{k',n}\left[\left(\boldsymbol{B}'_{i(n)}{}^{(t)}\left(\boldsymbol{B}'_{i(n)}{}^{(t)}\right)^T + \boldsymbol{\Lambda}_{i(n)}\right)^{-1}\right]_{k,k'}$$
$$+ \sum_{i=1}^{Q}\frac{1}{S_{\Delta Z_i}}\sum_{g=1}^{G}\sum_{k=1}^{K}\left(E_{\boldsymbol{\Delta}|\boldsymbol{Y},\Theta^{(t)}}\left(\Delta z_{i,g,k}\right) - (\boldsymbol{R}_k\boldsymbol{h}_i)_g\right)^2 + \sum_{i=1}^{Q}\frac{1}{S_{\Delta o_i}}\sum_{g=1}^{G}\left(E_{\boldsymbol{\Delta}|\boldsymbol{Y},\Theta^{(t)}}\left(\Delta o_{i,g}\right) - (\boldsymbol{R}_o\boldsymbol{h}_i)_g\right)^2$$
$$+ G\left(\sigma^{(t)}\right)^2\sum_{i=1}^{Q}\frac{1}{S_{\Delta Z_i}}\sum_{k=1}^{K}\left[\left(\boldsymbol{B}_i^{(t)}\left(\boldsymbol{B}_i^{(t)}\right)^T + \frac{1}{S_{\Delta Z_i}}\boldsymbol{I}\right)^{-1}\right]_{k,k} + G\left(\sigma^{(t)}\right)^2\sum_{i=1}^{Q}\frac{1}{S_{\Delta o_i}\left(N_i + \frac{1}{S_{\Delta o_i}}\right)}$$
$$+ \frac{1}{S_Z}\|\boldsymbol{Z}_r\|_F^2 + \frac{1}{S_o}\|\boldsymbol{o}_r - \mu_o\boldsymbol{1}_G\|_2^2$$

Combining Eq. (19)-(21), we have:

$$\widehat{\sigma^2} = \frac{\mathcal{S}}{GN + G + GQ + GK + GKQ + GL(K+1)} \tag{25}$$

###### 1.3.4 Solving other parameters

As Eq. (19)-(21) suggest, maximization of the expectation function with respect to all parameters other than $\boldsymbol{R}_o$, $\boldsymbol{R}_k$, and $\sigma^2$ can be done in the same way as in **sections 1.1.2-1.1.5**, with the exception that $\Delta\boldsymbol{Z}_i$ and $\Delta\boldsymbol{o}_i$ are replaced with their expectations, $E_{\Delta|\mathrm{Y},\Theta}(\Delta\boldsymbol{Z}_i)$ and $E_{\Delta|\mathrm{Y},\Theta}(\Delta\boldsymbol{o}_i)$, respectively.

##### 1.4 Fitting the GEDI model to UMI counts

In this section, we will discuss how the GEDI model can be fitted directly to the unnormalized, raw UMI count matrix $\boldsymbol{M}\in\mathbb{Z}^{G\times N}$. GEDI considers the following generative model for raw UMI counts:

$$m_{g,n} \sim \mathrm{Pois}(e^{y_{g,n}})$$
$$\boldsymbol{y}_n|\boldsymbol{\mu}_n(\Theta) \sim \mathcal{N}(\boldsymbol{\mu}_n(\Theta), \sigma^2\boldsymbol{I})$$
$$\Theta \sim g(\Theta)$$

Here, $\boldsymbol{\mu}_n(\Theta)$ is a function that returns the model-predicted vector $\boldsymbol{\mu}_n$ for cell $n$ given the model parameters $\Theta$ — for the case with no gene-level or sample-level prior information, this is defined as:

$$\boldsymbol{\mu}_n(\Theta) = \boldsymbol{o}_r + \Delta\boldsymbol{o}_{i(n)} + \left(\boldsymbol{Z}_r + \Delta\boldsymbol{Z}_{i(n)}\right)\boldsymbol{b}_n + s_n\boldsymbol{1}_G \tag{26}$$

When gene-level and sample-level prior information is given, the solution can be easily derived from the same concepts described in this section.

Here, the model parameter set includes:

$$\Theta = \left\{\boldsymbol{o}_r, \boldsymbol{Z}_r, \Delta\boldsymbol{o}_1, \ldots, \Delta\boldsymbol{o}_Q, \Delta\boldsymbol{Z}_1, \ldots, \Delta\boldsymbol{Z}_Q, \boldsymbol{B}, \sigma^2, \boldsymbol{s}\right\}$$

To obtain the MAP estimate for the parameters of this hierarchical model (in which $\boldsymbol{y}_n$ is a latent variable), we want to maximize the following density function:

$$\widehat{\Theta}_{\mathrm{MAP}}(\boldsymbol{M}) = \arg\max_\Theta P(\boldsymbol{M}|\Theta)g(\Theta) = \arg\max_\Theta \int P(\boldsymbol{M}|\Theta,\boldsymbol{Y})f(\boldsymbol{Y}|\Theta)d\boldsymbol{Y}\,g(\Theta)$$
$$= \arg\max_\Theta \int P(\boldsymbol{M}|\boldsymbol{Y})f(\boldsymbol{Y}|\Theta)d\boldsymbol{Y}\,g(\Theta)$$

We will achieve this by expectation maximization, where the expectation function $Q$ is:

$$Q\left(\Theta|\Theta^{(t)}\right) = E_{\boldsymbol{Y}|\boldsymbol{M},\Theta^{(t)}}[\log P(\boldsymbol{M}|\boldsymbol{Y})] + E_{\boldsymbol{Y}|\boldsymbol{M},\Theta^{(t)}}[\log f(\boldsymbol{Y}|\Theta)] + \log g(\Theta)$$

Note that, when maximizing the above function relative to $\Theta$, the first part can be ignored, which means that:

$$\arg\max_\Theta Q\left(\Theta\middle|\Theta^{(t)}\right) = \arg\max_\Theta \left[E_{\boldsymbol{Y}|\boldsymbol{M},\Theta^{(t)}}[\log f(\boldsymbol{Y}|\Theta)] + \log g(\Theta)\right]$$
$$= \arg\min_\Theta \left[-E_{\boldsymbol{Y}|\boldsymbol{M},\Theta^{(t)}}[\log f(\boldsymbol{Y}|\Theta)] - \log g(\Theta)\right]$$

Let's expand the first part:

$$-E_{\boldsymbol{Y}|\boldsymbol{M},\Theta^{(t)}}[\log P(\boldsymbol{Y}|\Theta)] = \frac{NG}{2} log(2\pi\sigma^2) + \frac{1}{2\sigma^2}\sum_{g=1}^{G}\sum_{n=1}^{N} E_{\boldsymbol{Y}|\boldsymbol{M},\Theta^{(t)}}\left[y_{g,n}^2 - 2y_{g,n}\mu_{g,n}(\Theta) + \mu_{g,n}^2(\Theta)\right]$$
$$= \frac{NG}{2} log(2\pi\sigma^2) + \frac{1}{2\sigma^2}\sum_{g=1}^{G}\sum_{n=1}^{N}\left[E_{\boldsymbol{Y}|\boldsymbol{M},\Theta^{(t)}}(y_{g,n}) - \mu_{g,n}(\Theta)\right]^2 + \frac{1}{2\sigma^2}\sum_{g=1}^{G}\sum_{n=1}^{N} Var_{\boldsymbol{Y}|\boldsymbol{M},\Theta^{(t)}}(y_{g,n})$$

Therefore:

$$\arg\max_\Theta Q\left(\Theta\middle|\Theta^{(t)}\right)$$
$$= \arg\min_\Theta \left[\frac{NG}{2} log(2\pi\sigma^2) + \frac{1}{2\sigma^2}\sum_{g=1}^{G}\sum_{n=1}^{N}\left[E_{\boldsymbol{Y}|\boldsymbol{M},\Theta^{(t)}}(y_{g,n}) - \mu_{g,n}(\Theta)\right]^2 - \log g(\Theta) + \frac{1}{2\sigma^2}\sum_{g=1}^{G}\sum_{n=1}^{N} Var_{\boldsymbol{Y}|\boldsymbol{M},\Theta^{(t)}}(y_{g,n})\right] \tag{27}$$

##### 1.4.1 Solving parameters other than $\sigma^2$

Eq. (27) suggests that all paramters can be estimated in the same way as explained in the **sections 1.1**, **1.2**, or **1.3** (depending on the presence of prior information), with the difference that $y_{g,n}$ should be replaced with its expectation given the UMI count $m_{g,n}$ and the current parameter estimates $\Theta^{(t)}$. Therefore, we will describe here how to obtain this expectation:

$$E_{\boldsymbol{Y}|\boldsymbol{M},\Theta^{(t)}}(y_{g,n}) = \int_{-\infty}^{+\infty} f\left(y_{g,n}\middle|m_{g,n}, \mu_{g,n}(\Theta)\right) y_{g,n} dy_{g,n}$$

For simplicity, in what follows, we drop the subscripts $g,n$ and simply write:

$$E_{\boldsymbol{Y}|\boldsymbol{M},\Theta^{(t)}}(y) = \int_{-\infty}^{+\infty} f\left(y\middle|m, \mu(\Theta)\right) y dy$$

Here, $f(y|m,\mu(\Theta))$ is the posterior probability density function given the observed count $m$ and the model estimate $\mu(\Theta)$, which can be expanded as follows:

$$f\left(y\middle|m, \mu(\Theta)\right) = \frac{P(m|y) f\left(y\middle|\mu(\Theta)\right)}{\int_{-\infty}^{+\infty} P(m|u) f\left(u\middle|\mu(\Theta)\right) du}$$
$$P(m|y) = \frac{(e^y)^m e^{-e^y}}{m!}$$
$$f\left(y\middle|\mu(\Theta)\right) = \frac{e^{-\frac{1}{2\sigma^2}\left(y-\mu(\Theta)\right)^2}}{\sigma\sqrt{2\pi}}$$

This leads to:

$$E_{\boldsymbol{Y}|\boldsymbol{M},\Theta^{(t)}}(y) = c\int_{-\infty}^{+\infty} (e^y)^m e^{-e^y} e^{-\frac{1}{2\sigma^2}\left(y-\mu(\Theta)\right)^2} y dy = c\int_{-\infty}^{+\infty} e^{ym - e^y - \frac{1}{2\sigma^2}\left(y-\mu(\Theta)\right)^2} y dy$$

where $c$ is a normalizing factor.

We use Laplace's method to approximate this integral, which leads to:

$$E_{\boldsymbol{Y}|\boldsymbol{M},\Theta^{(t)}}(y) \cong \arg\max_y \left[ym - e^y - \frac{1}{2\sigma^2}\left(y - \mu(\Theta)\right)^2\right] = \arg\max_y \left[y2\sigma^2 ym - 2\sigma^2 e^y - \left(y - \mu(\Theta)\right)^2\right]$$

By setting to zero the first derivative with respect to $y$, we have:

$$2\sigma^2 m - 2\sigma^2 e^y - 2\left(y - \mu(\Theta)\right) = 0$$
$$\sigma^2 e^y + y - \left(\mu(\Theta) + \sigma^2 m\right) = 0$$

$$y = -\mathrm{W}\left(\sigma^2 e^{\mu(\Theta)+\sigma^2 m}\right) + \mu(\Theta) + \sigma^2 m$$

Therefore:

$$E_{\boldsymbol{Y}|\boldsymbol{M},\Theta^{(t)}}(y) \cong= -W_0\left(\sigma^2 e^{\mu(\Theta)+\sigma^2 m}\right) + \mu(\Theta) + \sigma^2 m$$

where $W_0$ is the principal branch of the Lambert $W$ function. While this equation provides an exact solution, it is computationally expensive to calculate the Lambert $W$ function. Furthermore, when $\mu(\Theta)+\sigma^2 m$ is large, it is not possible to compute its exponential. Therefore, we use an alternative approach to approximate $y$ based on Halley's method.

Consider again the following equation:

$$h(y) = \sigma^2 e^y + y - (\mu(\Theta) + \sigma^2 m)$$

We can use Halley's method to update $y$ iteratively to find the root of $h$, using the equation:

$$y_{t+1} = y_t - \frac{2h(y_t)h'(y_t)}{2[h'(y_t)]^2 - h(y_t)h''(y_t)}$$
$$h'(y) = \sigma^2 e^y + 1$$
$$h''(y) = \sigma^2 e^y$$

In practice, we have found that this iterative procedure quickly converges on the solution for $y$. Note that calculating $h$ and its derivatives requires the calculation of $e^y$. Since $y$ is the logarithm of the expected read counts, $e^y$ is expected to remain in a range that is feasible to compute. However, it is also possible to rearrange the iterative equation as below:

$$y_{t+1} = y_t - \frac{2[e^{-y_t}h(y_t)][e^{-y_t}h'(y_t)]}{2[e^{-y_t}h'(y_t)]^2 - [e^{-y_t}h(y_t)][e^{-y_t}h''(y_t)]}$$
$$e^{-y_t}h(y_t) = \sigma^2 + e^{-y_t}(y_t - \mu(\Theta) - \sigma^2 m)$$
$$e^{-y_t}h'(y_t) = \sigma^2 + e^{-y_t}$$
$$e^{-y_t}h''(y_t) = \sigma^2$$

This provides the possibility of using the first set of equations when $y$ is negative, and the second set when $y$ is positive, in order to avoid calculation of the exponential of large positive numbers.

##### 1.4.2 Solving $\sigma^2$

Eq. (27) suggests that, to solve $\sigma^2$, the same equations as those presented in sections 1.1.6, 1.2.2, or 1.3.3 can be used (depending on the presence of prior information), with the only difference that the sum $\mathcal{S}$ should be modified to add the following term:

$$\mathcal{S}' = \mathcal{S} + \sum_{n=1}^{N} Var_{\boldsymbol{Y}|\boldsymbol{M},\Theta^{(t)}}(y_{j,n})$$

To obtain the variance of $y_{g,n}$ given the counts $m_{g,n}$ and the current model parameters, again we use the Laplace's approximation, where variance is estimated as the negative reciprocal of the second derivative of the $h$ function at its mode, where $h$ is defined as:

$$h(y) = ym - e^y - \frac{1}{2\sigma^2}\left(y - \mu(\Theta)\right)^2$$
$$h'(y) = m - e^y - \frac{1}{\sigma^2}\left(y - \mu(\Theta)\right)$$
$$h''(y) = -e^y - \frac{1}{\sigma^2} \Rightarrow -\frac{1}{h''(y)} = \frac{1}{e^y + \frac{1}{\sigma^2}}$$

Since the mode of the $h$ function is the expected value of $y$ (as described in **section 1.4.1**), we have:

$$Var_{\boldsymbol{Y}|\boldsymbol{M},\Theta^{(t)}}(y_{j,n}) = \frac{1}{e^{E_{\boldsymbol{Y}|\boldsymbol{M},\Theta^{(t)}}(y_{j,n})} + \frac{1}{\sigma^{2(t)}}}$$

#### 1.5 Fitting the GEDI model to paired UMI counts

In this section, we will discuss how the GEDI model can be fitted directly to a pair of unnormalized, raw UMI count matrices $\boldsymbol{M}_1 \in \mathbb{Z}^{G\times N}$ and $\boldsymbol{M}_2 \in \mathbb{Z}^{G\times N}$. We are interested in modeling the logarithm of odds ratio of observing a molecule from $\boldsymbol{M}_1$ vs. observing a molecule from $\boldsymbol{M}_2$. Accordingly, GEDI considers the following generative model for raw UMI counts:

$$m_{1,g,n} \sim B\left(m_{g,n}, \frac{1}{1+e^{-y_{g,n}}}\right)$$
$$m_{g,n} = m_{1,g,n} + m_{2,g,n}$$
$$\boldsymbol{y}_n | \boldsymbol{\mu}_n(\Theta) \sim \mathcal{N}(\boldsymbol{\mu}_n, \sigma^2 \boldsymbol{I})$$
$$\Theta \sim g(\Theta)$$

Here, $\boldsymbol{\mu}_n(\Theta)$ is similar to **section 1.4**, Eq. (26); each element $\mu_{n,g}$ represents the model-predicted log odds ratio of $m_{1,n,g}$ vs. $m_{2,n,g}$. $B$ is the binomial distribution.

Following the same procedure as in **section 1.4**, we can see that, for estimation of the model parameters using expectation-maximization, we need to maximize the following function:

$$\arg\max_\Theta Q\left(\Theta \middle| \Theta^{(t)}\right)$$
$$= \arg\min_\Theta \left[\frac{NG}{2} log(2\pi\sigma^2) + \frac{1}{2\sigma^2}\sum_{g=1}^{G}\sum_{n=1}^{N}\left[E_{\boldsymbol{Y}|\boldsymbol{M}_1,\boldsymbol{M},\Theta^{(t)}}(y_{g,n}) - \mu_{g,n}(\Theta)\right]^2 - \log g(\Theta) + \frac{1}{2\sigma^2}\sum_{g=1}^{G}\sum_{n=1}^{N} Var_{\boldsymbol{Y}|\boldsymbol{M}_1,\boldsymbol{M},\Theta^{(t)}}(y_{g,n})\right]$$

Therefore, the same concepts as those outlined in **sections 1.4.1** and **1.4.2** apply here, except that we need to calculate $E_{\boldsymbol{Y}|\boldsymbol{M}_1,\boldsymbol{M},\Theta(t)}(y_{g,n})$ and $Var_{\boldsymbol{Y}|\boldsymbol{M}_1,\boldsymbol{M},\Theta(t)}(y_{g,n})$.

##### 1.5.1 Obtaining the expectation of *Y*

Similar to section 1.4.1, we have:

$$E_{\boldsymbol{Y}|\boldsymbol{M}_1,\boldsymbol{M},\Theta^{(t)}}(y_{g,n}) = \int_{-\infty}^{+\infty} f\left(y_{g,n}\middle| m_{1,g,n}, m_{g,n}, \mu_{g,n}(\Theta)\right) y_{g,n} dy_{g,n}$$

For simplicity, in what follows, we drop the subscripts $g,n$ and simply write:

$$E_{\boldsymbol{Y}|\boldsymbol{M}_1,\boldsymbol{M},\Theta^{(t)}}(y) = \int_{-\infty}^{+\infty} f\big(y\big|m_1, m, \mu(\Theta)\big) y dy$$

Here, $f(y|m_1,m,\mu(\Theta))$ is the posterior probability density function given the observed counts $m_1$ and $m=m_1+m_2$ and the model estimate $\mu(\Theta)$:

$$f\big(y\big|m_1, m, \mu(\Theta)\big) = \frac{P(m_1|m,y) f\big(y\big|\mu(\Theta)\big)}{\int_{-\infty}^{+\infty} P(m_1|m,u) f\big(u\big|\mu(\Theta)\big) du}$$
$$P(m_1|m,y) = \binom{m}{m_1}\left(\frac{1}{1+e^{-y}}\right)^{m_1}\left(\frac{1}{1+e^{y}}\right)^{m-m_1}$$
$$f\big(y\big|\mu(\Theta)\big) = \frac{e^{-\frac{1}{2\sigma^2}\left(y-\mu(\Theta)\right)^2}}{\sigma\sqrt{2\pi}}$$

This leads to:

$$E_{\boldsymbol{Y}|\boldsymbol{M}_1,\boldsymbol{M},\Theta^{(t)}}(y) = c\int_{-\infty}^{+\infty}\left(\frac{1}{1+e^{-y}}\right)^{m_1}\left(\frac{1}{1+e^{y}}\right)^{m-m_1} e^{-\frac{1}{2\sigma^2}\left(y-\mu(\Theta)\right)^2} y dy =$$
$$= c\int_{-\infty}^{+\infty} (e^y)^{m_1}(1+e^y)^{-m} e^{-\frac{1}{2\sigma^2}\left(y-\mu(\Theta)\right)^2} y dy$$

where $c$ is a normalizing factor.

We use Laplace's method to approximate this integral, which leads to:

$$E_{\boldsymbol{Y}|\boldsymbol{M}_1,\boldsymbol{M},\Theta^{(t)}}(y) \cong \arg\max_y\left\{m_1 y - m\ln(1+e^y) - \frac{1}{2\sigma^2}\left(y-\mu(\Theta)\right)^2\right\}$$

(Note that the function that is to be maximized is concave, since its second derivative with respect to $y$ is always negative).

By setting to zero the first derivative with respect to $y$, we have:

$$\frac{d}{dy}\left[m_1 y - m\ln(1+e^y) - \frac{1}{2\sigma^2}\big(y-\mu(\Theta)\big)^2\right] = m_1 - \frac{m}{1+e^{-y}} - \frac{y-\mu(\Theta)}{\sigma^2} = 0$$

With some rearrangements, this leads to:

$$\sigma^2 m_2 - \mu(\Theta) + y + (y - \mu(\Theta) - \sigma^2 m_1)e^{-y} = 0$$

This can be rewritten as:

$$h(y) = y - \alpha + (y-\beta)e^{-y} = 0$$

where $\alpha$ and $\beta$ are defined as:

$$\alpha = \mu(\Theta) - \sigma^2 m_2$$
$$\beta = \mu(\Theta) + \sigma^2 m_1$$

We will find the root of the function $h$ (above) using Halley's method:

$$h(y) = y - \alpha + (y-\beta)e^{-y}$$
$$h'(y) = 1 + e^{-y}(\beta - y + 1)$$
$$h''(y) = e^{-y}(y - \beta - 2)$$
$$y_{t+1} = y_t - \frac{2h(y_t)h'(y_t)}{2[h'(y_t)]^2 - h(y_t)h''(y_t)}$$

Note that when $-y$ is large, the solution above requires computation of $e^{-y}$, which can result in numerical instability or out of range results. We can instead use the following calculations:

$$y_{t+1} = y_t - \frac{2[e^{y_t}h(y_t)][e^{y_t}h'(y_t)]}{2[e^{y_t}h'(y_t)]^2 - [e^{y_t}h(y_t)][e^{y_t}h''(y_t)]}$$
$$e^{y_t}h(y_t) = e^{y_t}(y_t - \alpha) + y_{nt} - \beta$$
$$e^{y_t}h'(y_t) = e^{y_t} + \beta - y_t + 1$$
$$e^{y_t}h''(y_t) = y_t - \beta - 2$$

In practice, we have found that Halley's method may overshoot or undershoot in some cases. We resolve this problem by identifying an upper and lower bound for the root of function $h$, within which Halley's method results in convergence to the solution. It is easy to show that the root of $h$ must be between $\alpha$ and $\beta$ ($\alpha<\beta$) because $h(\alpha)<0$ and $h(\beta)>0$. Furthermore, within this range, $h'(y)>0$ (i.e. $h$ is monotonically increasing), and $h''(y)<0$, meaning that Halley's method will not over/undershoot if initialized with $\alpha$.

##### 1.5.2 Obtaining the variance of *Y*

To obtain the variance of $y_{g,n}$ given the counts $m_{1,g,n}$ and $m_{2,g,n}$ and the current model parameters, again we use the Laplace's approximation, where variance is estimated as the negative reciprocal of the second derivative of the $h$ function at its mode, where $h$ is defined as:

$$h(y) = m_1 y - m\ln(1+e^y) - \frac{1}{2\sigma^2}\big(y-\mu(\Theta)\big)^2$$
$$h'(y) = m_1 - \frac{m}{1+e^{-y}} - \frac{y-\mu(\Theta)}{\sigma^2}$$
$$h''(y) = -\frac{me^{-y}}{(1+e^{-y})^2} - \frac{1}{\sigma^2}$$
$$-\frac{1}{h''(y)} = \frac{1}{\frac{me^{-y}}{(1+e^{-y})^2} + \frac{1}{\sigma^2}} = \frac{1}{\frac{me^{y}}{(1+e^{y})^2} + \frac{1}{\sigma^2}}$$

And since mode of the $h$ function is the expected value of $y$, we have:

$$Var_{\boldsymbol{Y}|\boldsymbol{M_1},\boldsymbol{M},\Theta^{(t)}}(y_{g,n}) = \frac{1}{\frac{me^{-\left|E_{\boldsymbol{Y}|\boldsymbol{M_1},\boldsymbol{M},\Theta^{(t)}}(y)\right|}}{\left(1+e^{-\left|E_{\boldsymbol{Y}|\boldsymbol{M_1},\boldsymbol{M},\Theta^{(t)}}(y)\right|}\right)^2} + \frac{1}{\sigma^2_{(t)}}}$$

#### 1.6 Choice of hyperparameters

The behavior of GEDI may be fine-tuned using its hyperparameters, which primarily represent the variances of the prior distributions of the model parameters. However, in the present work, we have not systematically explored the effects of increasing or decreasing various prior distribution variances, as well as potential metrics that may be used to fine-tune the hyperparameters for a given dataset. Instead, we have selected a set of predefined values based on what we would expect to infer from "random" gene expression data, i.e., data generated from a model in which all parameters are zero except for the model variance $\sigma^2$, as described below.

##### 1.6.1 The prior distribution of $Z_r$

Consider the matrix of observed gene expression $\boldsymbol{Y}$ generated from a model similar to that described in **section 1.1**, with the "null" parameter set $\Theta_0$ in which all parameters, except $\sigma^2$, are zero. We then use $\boldsymbol{Y}$ to infer $\Theta$ using a block coordinate descent approach similar to that used by GEDI, with the exception that we use flat priors for each parameter. When inferring $\boldsymbol{Z}_r$ with the flat prior U(–∞,+∞), each row $\boldsymbol{z}_{r,g,*}$ of $\boldsymbol{Z}_r$ will have the following posterior distribution, conditional on $\boldsymbol{Y}$ and all other model parameter estimates:

$$\boldsymbol{z}_{r,g,*}|\boldsymbol{Y},\widehat{\Theta}\backslash\{\widehat{\boldsymbol{Z}}_r\}\sim\mathcal{N}\left(\hat{\boldsymbol{z}}_{r,g,*},\hat{\sigma}^2\left(\widehat{\boldsymbol{B}}\widehat{\boldsymbol{B}}^\top\right)^{-1}\right)$$

Here, $\hat{\boldsymbol{z}}_{r,g,*}$ is the ML estimate of $\boldsymbol{z}_{r,g,*}$ given $\boldsymbol{Y}$ and other model parameters. Note that $\boldsymbol{B}$ is restricted to have rows with $L^2$ norm of 1, as described in **section 1.1.5**. Furthermore, under the assumptions of the null model, the expected covariance of different rows of $\boldsymbol{B}$ is zero, leading to $E_{\boldsymbol{Y}|\Theta_0}(\hat{\boldsymbol{B}}\hat{\boldsymbol{B}}^\top)=\boldsymbol{I}$. Therefore:

$$E_{\boldsymbol{Y}|\Theta_0}\left(Cov_{\boldsymbol{z}_{r,g,*}|\boldsymbol{Y},\widehat{\Theta}\backslash\{\hat{\boldsymbol{z}}_r\}}\left(\boldsymbol{z}_{r,g,*}\right)\right)=\hat{\sigma}^2\boldsymbol{I}$$

In other words, each element of $z_{r,g,k}$ is expected to have a posterior normal distribution with mean $\hat{z}_{r,g,k}$ and variance $\sigma^2$. Conversely, we can infer that the ML estimate $\hat{z}_{r,g,k}$ is expected to be sampled from a normal distribution with mean 0 and variance $\sigma^2$. We use this distribution as the prior in the GEDI model:

$$\boldsymbol{z}_{r,*,k}\sim\mathcal{N}(\boldsymbol{0},\sigma^2 S_Z\boldsymbol{I})$$

where $S_Z$ is set to 1.

##### 1.6.2 The prior distribution of $\Delta Z_i$

Let's consider the same null model as the previous section. When inferring $\Delta\boldsymbol{Z}_i$ (for each $i\in\{1,\ldots,Q\}$) from data generated by this null model, each row $\Delta\boldsymbol{z}_{i,g,*}$ of $\Delta\boldsymbol{Z}_i$ will have the following posterior distribution if a flat prior U(–∞,+∞) is used, conditional on $\boldsymbol{Y}$ and all other model parameter estimates:

$$\Delta\boldsymbol{z}_{i,g,*}|\boldsymbol{Y},\widehat{\Theta}\backslash\{\Delta\widehat{\boldsymbol{Z}}_i\}\sim\mathcal{N}\left(\Delta\hat{\boldsymbol{z}}_{i,g,*},\hat{\sigma}^2\left(\widehat{\boldsymbol{B}}_i\widehat{\boldsymbol{B}}_i^\top\right)^{-1}\right)$$

Following the same assumptions as those of the previous section, we can see that $E_{\boldsymbol{Y}|\Theta_0}(\hat{\boldsymbol{B}}_i\hat{\boldsymbol{B}}_i^\top)=(N_i/N)\times\boldsymbol{I}$, where $N_i$ is the number of cells in sample $i$ and $N$ is the total number of cells across all samples. This suggests the following prior:

$$\Delta\boldsymbol{z}_{i,*,k}\sim\mathcal{N}\left(\boldsymbol{0},\sigma^2 S_{\Delta Z_i}\boldsymbol{I}\right)$$

$$S_{\Delta Z_i}=\frac{N}{N_i}$$

##### 1.6.3 The prior distribution of $\Delta o_i$

Similar to the previous sections, we can see that, with a flat prior, the posterior distribution of each element of the $\Delta\boldsymbol{o}_i$ vector, when $\boldsymbol{Y}$ is generated from a null parameter set, is given by:

$$\Delta o_{i,g}|\boldsymbol{Y},\widehat{\Theta}\backslash\{\Delta\hat{\boldsymbol{o}}_i\}\sim\mathcal{N}\left(\Delta\hat{o}_{i,g},\hat{\sigma}^2\frac{1}{N_i}\right)$$

Therefore, we use the following prior:

$$\Delta\boldsymbol{o}_i\sim\mathcal{N}\left(\boldsymbol{0},\sigma^2 S_{\Delta o_i}\boldsymbol{I}\right)$$

$$S_{\Delta o_i}=\frac{1}{N_i}$$

We note that $\Delta\boldsymbol{o}_i$ represents a simple linear batch correction that maps the centroids of the manifolds of the different samples to the centroid of the reference manifold. Therefore, when all samples are expected to have similar cell types, a less stringent prior can be placed on $\Delta\boldsymbol{o}_i$. In the benchmarking analyses presented in this paper, we use $S_{\Delta oi}=1000/N_i$ to better accommodated the mean abundance differences that exist between multiple scRNA-seq technologies.

##### 1.6.4 The prior distribution of $A$

In the presence of gene-level prior information (represented by matrix $\boldsymbol{C}$), $\boldsymbol{Z}_r$ is modeled as:

$$\boldsymbol{z}_{r,k}|\boldsymbol{a}_k\sim\mathcal{N}(\boldsymbol{C}\boldsymbol{a}_k,\sigma^2 S_Z\boldsymbol{I})$$

Similar to the previous sections, we consider data generated from a null model in which $\boldsymbol{A}=\boldsymbol{0}$. When inferring $\boldsymbol{A}$ from this data with a flat prior, conditional on $\boldsymbol{Z}_r$ and $\sigma^2$, the posterior distribution of $\boldsymbol{A}$ is given by:

$$\boldsymbol{a}_k \big| \widehat{\boldsymbol{Z}}_r, \hat{\sigma}^2 \sim \mathcal{N}(\widehat{\boldsymbol{a}}_k, \hat{\sigma}^2 S_Z (\boldsymbol{C}^\top \boldsymbol{C})^{-1})$$

If $\boldsymbol{C}$ has orthonormal column vectors, then $\boldsymbol{C}^\top\boldsymbol{C}=\boldsymbol{I}$, and since $S_z=1$, the posterior distribution of each element of $\boldsymbol{a}_k$ will have a variance equal to $\sigma^2$. We can therefore use the following prior:

$$\boldsymbol{a}_k \sim \mathcal{N}(\boldsymbol{0}, \sigma^2 S_A \boldsymbol{I})$$
$$S_A = 1$$

GEDI uses this prior even when the provided matrix $\boldsymbol{C}$ does not have orthonormal column vectors. However, in this case, it uses singular value decomposition (SVD) to obtain $\boldsymbol{C}=\boldsymbol{U\Sigma V}^\top$. It then uses the matrix $\boldsymbol{C}'=\boldsymbol{U}$ instead of $\boldsymbol{C}$ in model fitting. Once the matrix $\boldsymbol{A}'$ is obtained by fitting the model with $\boldsymbol{C}'$ instead of $\boldsymbol{C}$, $\boldsymbol{A}$ can be obtained as follows:

$$\boldsymbol{CA} = \boldsymbol{C}'\boldsymbol{A}'$$
$$\Rightarrow \boldsymbol{C}'\boldsymbol{\Sigma V}^\top \boldsymbol{A} = \boldsymbol{C}'\boldsymbol{A}'$$
$$\Rightarrow \boldsymbol{\Sigma V}^\top \boldsymbol{A} = \boldsymbol{A}'$$
$$\Rightarrow \boldsymbol{A} = \boldsymbol{V\Sigma}^{-1}\boldsymbol{A}'$$

Note that this is equivalent to a regularized principal component regression.

##### 1.6.5 The prior distributions of $\boldsymbol{R}_o$ and $\boldsymbol{R}_k$

In the presence of sample-level prior information (represented by vector $\boldsymbol{h}_i$ for each sample $i$), $\Delta\boldsymbol{o}_i$ is modeled as:

$$\Delta\boldsymbol{o}_i | \boldsymbol{R}_o \sim \mathcal{N}\left(\boldsymbol{R}_o \boldsymbol{h}_i, \sigma^2 S_{\Delta o_i} \boldsymbol{I}\right)$$

If we infer $\boldsymbol{R}_o$ with a flat prior, conditional on $\Delta\boldsymbol{O}$ (as defined in **section 1.3.1**) and $\sigma^2$ we have:

$$\boldsymbol{r}_{o,g,*} \big| \Delta\widehat{\boldsymbol{O}}, \hat{\sigma}^2 \sim \mathcal{N}\left(\hat{\boldsymbol{r}}_{o,g,*}, \hat{\sigma}^2 \left(\boldsymbol{H}'_{\Delta \boldsymbol{O}} \boldsymbol{H}'^{\top}_{\Delta \boldsymbol{O}}\right)^{-1}\right)$$

where $\boldsymbol{r}_{o,g,*}$ represents row $g$ of matrix $\boldsymbol{R}_o$, and, similar to **section 1.3.1**, $\boldsymbol{H}'_{\Delta \boldsymbol{O}}$ is defined as:

$$\boldsymbol{H}'_{\Delta \boldsymbol{O}} = \left[\frac{1}{\sqrt{S_{\Delta o_1}}}\boldsymbol{h}_1 \quad \dots \quad \frac{1}{\sqrt{S_{\Delta o_Q}}}\boldsymbol{h}_Q\right]$$

If $\boldsymbol{H}'_{\Delta \boldsymbol{O}}$ has orthonormal row vectors, then similar to the previous section, we can see that the covariance of the posterior distribution of each row of $\boldsymbol{R}_o$ will be $\sigma^2\boldsymbol{I}$, which leads us to use the following prior:

$$\boldsymbol{R}_o \sim \mathcal{N}\left(\boldsymbol{0}, \sigma^2 S_{R_o} \boldsymbol{I}\right)$$
$$S_{R_o} = 1$$

A similar line of calculations will lead to the following prior matrix for Rk as long as $\boldsymbol{H}'_{\Delta \boldsymbol{Z}}$ has orthonormal rows:

$$\boldsymbol{R}_k \sim \mathcal{N}\left(\boldsymbol{0}, \sigma^2 S_{R_k} \boldsymbol{I}\right)$$
$$S_{R_k} = 1$$

GEDI uses SVD to convert $\boldsymbol{H}'_{\Delta \boldsymbol{O}}$ and $\boldsymbol{H}'_{\Delta \boldsymbol{Z}}$ to matrices with orthonormal rows, allowing it to use the above priors. The fitted matrices $\boldsymbol{R}'_o$ and $\boldsymbol{R}'_k$ are then converted to the original scale similar to the previous section.

#### 2 Datasets and preprocessing

##### 2.1.1 PBMC dataset

The PBMC dataset[1] is a collection of immune cell types profiled from peripheral blood from two human donors across six single-cell RNA-seq technologies (10x Chromimum v2 and v3, CEL-seq2, Drop-seq, inDrops, Seq-Well and Smart-seq2). The raw count matrices and metadata were retrieved using the SeuratData package (v0.1), using the accession code "pbmcsca" (v.3.0.0). The data can be downloaded at the original website repository[2] or at the GEO accession number GSE132044.

For QC, we kept cells that passed the following cutoffs: $\log_{10}$ total counts >2.5, $\log_{10}$ total features >2, and percentage of mitochondrial reads per cell <20%. To remove low-expressed genes, we kept genes that had more than five counts in more than three cells across the entire dataset. After filtering, the PBMC dataset contained 28,035 cells and 14,406 genes. To obtain normalized expression counts, we used the function multiBatchNorm from batchelor (v.1.6.3). For the integration benchmark, we used the combination of the technology and the donor variable as the 'batch' variable (14 samples).

##### 2.1.2 Pancreas dataset

The Pancreas dataset is a collection of datasets from multiple studies that profiled human cells from the pancreas using various scRNAseq technologies. The raw count matrices and metadata were retrieved using the SeuratData package[3] (v0.1), using the accession code "panc8" (v.3.0.2). The original collection consisted of eight datasets (celseq, celseq2, fluidigmc1, indrop1, indrop2, indrop3, indrop4 and smartseq2). As we were interested in comparing the performance of GEDI when fitted directly to raw counts against the performance of other methods, we restricted the analysis to the datasets that contained raw count data available (indrop1, indrop2, indrop3, indrop4 and smartseq2). The Smart-seq2 dataset[4] contained five healthy donors and four donors with type 2 diabetes—the original processed data can be found at ArrayExpress accession number E-MTAB-5061. The inDrop datasets[5] contained four healthy donors, and the original processed data can be found at GEO accession number GSE84133.

For QC, we kept cells that passed the following cutoffs: $\log_{10}$ total counts >3 and $\log_{10}$ total features >2.8. To alleviate the effect of donors with low cell numbers, we removed samples that had less than 100 cells per donor. To remove low-expressed genes, we kept genes that had more than five counts in more than three cells across the entire dataset. After filtering, the Pancreas dataset contained 10,902 cells and 18,366 genes. To obtain normalized expression counts, we used the function multiBatchNorm from batchelor[6, 7] (v.1.6.3). For the integration benchmark, we used the donor (encoded as 'orig.ident' in the metadata) as the 'batch' variable (13 samples).

##### 2.1.3 Tabula Muris BM dataset

The Tabula Muris BM dataset[8] is a collection of cell types profiled from the bone marrow of mice across two scRNA-seq technologies (10x Chromimum and Smart-seq2). Data in H5AD format for the 10x experiment was retrieved from ref[9]. We restricted our analysis to bone marrow cells derived from three female mice.

For QC, we kept cells that passed the following cutoffs: $\log_{10}$ total counts >3 and $\log_{10}$ total features >3. To remove low-expressed genes, we kept genes that had more than five counts in more than three cells across the entire dataset. After filtering, the Tabula Muris BM dataset contained 13,874 cells and 15,472 genes. To obtain normalized expression counts, we used the function multiBatchNorm from batchelor (v.1.6.3). For the integration benchmark, as the mice of origin were different across both scRNA-seq technologies, we used the mouse of origin (encoded as 'mouse.id' in the metadata) as the 'batch' variable (10 samples).

##### 2.1.4 COVID-19 dataset

The COVID-19 dataset[10] profiled human peripheral blood samples from independent patient cohorts at two university medical centers in Germany. Samples from cohort 1 were profiled using 10x Chromium, while samples from cohort 2 were profiled using a microwell-based scRNA-seq system (Rhapsody). The dataset contains individuals diagnosed with mild and severe COVID-19, as well as healthy controls. Raw count data and metadata were retrieved from the processed Seurat objects, which were downloaded from the FastGenomics Portal; Cohort 1 dataset was downloaded from[11], while Cohort 2 dataset was downloaded from[12]. The data is deposited at the European Genome-phenome Archive (EGA) under access number EGAS00001004571.

For QC, we kept cells that passed the following cutoffs: $\log_{10}$ total counts >3, $\log_{10}$ total features >2, and percentage of mitochondrial reads per cell <20%. To remove low-expressed genes, we kept genes that had more than three counts in more than three cells across the entire dataset. After filtering, the COVID-19 dataset contained 197,039 cells and 13,205 genes. To obtain normalized expression counts, we used the function multiBatchNorm from batchelor (v.1.6.3). For the analysis of sample-to-sample variability with GEDI, the donor of origin was considered as the 'sample' variable, except for donors BN-10, BN-11 and BN-12—each of those three patients were profiled in severe and mild conditions, so a combination of the donor and the COVID status was used.

##### 2.1.5 Tasic dataset

The Tasic dataset is a collection of two studies that profiled neocortex tissue in adult mice. The Tasic 2016 study[13] used the SMARTer Ultra Low RNA Kit and generated single-end reads, while the Tasic 2018 study[14] used the SMART-Seq v4 Ultra Low Input RNA Kit for Sequencing protocol and generated paired-end reads. FASTQ files were downloaded from the NCBI Short Read Archive (SRA) under accession numbers SRP061902 and SRP150473. Metadata for the Tasic 2016 dataset was retrieved from the GEO accession number GSE71585, as well as from the Supplementary Table 1 from Feng et al.[15], who performed a splicing-analysis of the two Tasic datasets. Metadata for the Tasic 2018 dataset was retrieved from the GEO accession number GSE115746.

Quantification of the reads that support exon inclusion and exon exclusion events was performed using the Quantas pipeline[16] (v.1.1.1). Alignment of the reads to the mm10 genome was performed using Olego[17, 18] (v.1.1.9). To quantify alternative splicing events, we inferred the transcript structure between paired-end reads using 'gapless' (this step was only performed for the paired-end data). Then, we quantified inclusion or exclusion read counts for cassette exons using the 'summarize_splicing_wrapper.pl' script from 'countit'.

For QC, we filtered out cells that were classified as low-quality cells in the original metadata. Then, we restricted our analysis to cells that were classified as 'Non-Neuronal', 'GABAergic', 'Glutamatergic', and 'Endothelial' in the original metadata. Finally, we kept cells that passed the following cutoff: $\log_{10}$ total inclusion counts >4.5 and $\log_{10}$ total exclusion counts >4. To remove low-expressed cassette exons, we kept events that had more than 50 exon inclusion counts and more than 50 exon exclusion counts across the entire dataset. After filtering, the Tasic dataset contained 25,352 cells and 14,267 exon events. Normalized exclusion and inclusion counts were obtained using the normalizeCounts from scuttle.

For the integration tasks, for all methods except for GEDI-B and LIGER, we used the logarithm of a naïve estimate of the ratio of the inclusion and exclusion counts as input for each method. This ratio was calculated as $(1+M'_1)/(1+M'_2)$, where $M'_1$ represents the normalized count matrix for the inclusion counts and $M'_2$ represents the normalized count matrix for the exclusion counts. For LIGER, we used $(1+M_1)/(2+M_1+M_2)$, where $M_1$ represents the raw count matrix for the inclusion counts and $M_2$ represents the raw count matrix for the exclusion counts. These choices were made based on the requirement for the input data of each method; for example, while most other methods can work with values that span negative and positive numbers, LIGER works with only positive numbers. For GEDI-B (GEDI with a binomial data generating distribution), we used the pair of raw inclusion and exclusion counts as input.

##### 2.1.6 Faure dataset

Faure et al.[19] used scRNA-seq (Smart-seq2) to understand the developmental diversity that occurs in sensory neurogenesis in mice. The processed data was downloaded in LOOM format from the GEO accession number GSE150150, which contained intronic and exonic raw counts. Metadata was downloaded from the GitHub repository[20] associated with the article. Only cells that passed QC in the original article were used for analysis. To remove low-expressed genes, we kept genes that (a) had more than five counts in more than three cells across the entire dataset, (b) were expressed in more than one cell in the intronic counts, and (c) were expressed in more than one cell in the exonic counts. After filtering, the Faure dataset consisted of 2,245 cells and 14,725 genes.

##### 2.1.7 La Manno dataset

La Manno et al.[21] used scRNA-seq (10x Chromium) to understand the kinetics of transcription in human embryonic glutamatergic neurogenesis. The processed data was downloaded in LOOM format from the authors' repository[22], which contained metadata, intronic and exonic counts. Only cells that passed QC in the original article were used for analysis. To remove low-expressed genes, we kept genes that (a) had more than five counts in more than three cells across the entire dataset, (b) were expressed in more than one cell in the intronic counts, and (c) were expressed in more than one cell in the exonic counts. After filtering, the La Manno dataset consisted of 1,720 cells and 5,037 genes.

#### 3 Integration methods

To compare the integration performance across methods, we ran each method using the same set of genes (all genes after filtering low-expressed genes) and five different values of $K$ (where $K$ represents the number of latent variables), including 20, 40, 60, 80, and 100. The details of each method can be found below:

##### 3.1.1 GEDI

For the integration benchmark tasks (PBMC, Pancreas, and Tabula Muris BM), we ran GEDI using the raw counts and with default parameters, except that we set 'oi_shrinkage=0.001'. The integrated embedding was retrieved using the function 'svg.gedi'. For non-benchmark integration tasks, $K$=20 was used.

##### 3.1.2 Seurat

To run Seurat[23, 24] (v.4.1.1), we followed the documentation available on the Seurat website[25]. We ran the Canonical Correlation Analysis (CCA) pipeline for integration in Seurat, which included the functions CreateSeuratObject, SplitObject(split.by=batch), NormalizeData, FindIntegrationAnchors and IntegrateData. To run Seurat with a specified gene list, these were indicated in the FindIntegrationAnchors with the 'anchor.features' parameter. PCA was computed on the integrated expression matrix returned by Seurat.

##### 3.1.3 LIGER

To run LIGER[26] (v.1.0.0), we followed the documentation available on the LIGER GitHub repository[27]. We applied the pipeline used for the integration of multiple scRNA-seq datasets, which performs integrative nonnegative matrix factorization (iNMF). This included the createLiger, normalize, scaleNotCenter, optimizeALS and quantile_norm functions. The integrated embedding returned by LIGER was obtained from the 'H.norm' slot space in the LIGER object.

##### 3.1.4 Harmony

To run Harmony[28, 29] (v.1.0), we followed the documentation available on the Harmony GitHub repository[30]. We first ran PCA on the normalized expression matrix of each dataset and then used the HarmonyMatrix function with arguments 'do_pca=FALSE' to obtain the integrated Harmony embeddings.

##### 3.1.5 BBKNN

To run BBKNN[31] (v.1.3.12), we followed the integration tutorial on the GitHub repository[32], which suggested applying the pre-processing steps with scanpy[33] (v.1.9.1) and then performing integration with BBKNN. We ran the normalization steps (scanpy.pp.normalize_per_data and scanpy.pp.log1p) and computed PCA using scanpy.tl.pca. The integrated space was recovered using the bbknn.bbknn function, which constructs a batch-balanced neighborhood graph.

##### 3.1.6 CSS

To run CSS[34] (simspec v.0.0.0.9000), we followed the documentation available in the GitHub repository[35]. CSS uses a processed Seurat Object, so we first created a Seurat object and followed the default scRNA-seq pipeline (CreateSeuratObject, NormalizeData, ScaleData, RunPCA). Then, we used the cluster_sim_spectrum from simspec, which returned CSS-integrated embeddings.

##### 3.1.7 PCA

PCA was run using the rpca function from rsvd[36] (v.1.0.5) on the normalized expressed data.

#### 4 Metrics to compare integration performance

All metrics described below rely on identification of the neighborhood of each cell, followed by examination of the extent to which cells from different batches or different cell types mix together within the neighborhoods. It is desired to see the cells from different batches mix together within neighborhoods, while cells from different cell types should remain in separate neighborhoods. Identification of the neighborhoods relies on measuring the pairwise distances of the cells in the integrated space. Given that the integrated space is still high-dimensional (20-100 dimensions) and, therefore, Euclidean distances between pairs of points may become relatively homogenous (curse of dimensionality), we first generated a UMAP[37] embedding of the cells with the same number of dimensions as that of the original integrated space ($K$). The UMAP transformation leads to an emphasis on the local connectivity of the points. Furthermore, as previously shown[38], Euclidean distances of points in the UMAP embedding better correlate with their geodesic distances, especially when the data points are noisy, compared to Euclidean distances in, for example, PCA-projected data.

For all methods except BBKNN, the output of each method was an integrated embedding, from which we generated a UMAP embedding of dimension $K$, using the umap function from uwot[39] (v.0.1.10). For BBKNN, we ran UMAP on the batch-balanced neighborhood graph. The K-dimensional UMAP embedding was used as input for the calculation of the integration metrics, as described below.

##### 4.1.1 Alignment Score

The alignment score was proposed by Butler et al.[40], with the goal of quantifying how well any group of data sets is aligned. The alignment score ranges from 0 to 1, and when applied to batch labels, a value closer to 1 denotes good mixing. We implemented the alignment score as an R function as defined previously[40]. The alignment score builds a k-nearest neighbor graph based on the cell's embedding; we used k=10. The alignment score was applied either to the cell type or batch labels. For the cell type labels, to ensure that a higher AS score means better cell type conservation, we subtracted it from 1 to give a final cell type AS score.

##### 4.1.2 LISI

The local inverse Simpson's Index (LISI) was proposed by Korsunsky et al.[29], and consists of a measure to assess batch mixing (integration LISI) and cell-type separation (cell-type LISI). Integration LISI (iLISI) defines the effective number of batches in a neighborhood, and a score close to the expected number of batches denotes good mixing. We used the compute_lisi function from lisi[41] (v.1.0) on the cell type labels (cLISI) or on the batch labels (iLISI). The original LISI scores range from 1 to $B$ (where $B$ is the number of batches), so we applied the normalization procedure proposed by Luecken et al.[42], where the values of LISI were rescaled to range from 0 to 1. Specifically, scaled iLISI was calculated as [median(iLISI)–1]/[$B$–1], and scaled cLISI was calculated as [$B$–median(cLISI)]/[$B$–1].

##### 4.1.3 kBET

kBET was developed by Büttner et al.[43] as a measure to quantify batch effects in scRNA-seq data. kBET uses a chi-squared test to assess the mixing of fixed-size random neighborhoods. An overall rejection rate is calculated after averaging the binary test results, with low rejection rates indicating well-mixed batches. We used the kBET function from kBET package[44] (v0.99.6), using 'k0=30' and 'do.pca=FALSE' on the batch labels. To ensure that a higher kBET score represents better batch removal, we subtracted the original score from 1 to give a final kBET score.

##### 4.1.4 ASW

The silhouette width measures the relationship between the similarity of a point to its own cluster and the similarity of that point to the closest neighboring cluster. The score ranges from -1 to 1, where a score close to 1 represents that the point is properly clustered, while a score $\leqslant 0$ represents poor correspondence to its own cluster. The average silhouette width (ASW) provides an evaluation of clustering validity. To compute ASW, we used the 'silhouette' function from the 'cluster' R package[45] (v.2.1.0). To compute cell type ASW and batch ASW, we followed the scaling and normalization established previously[42]. Specifically, cell type ASW ($\mathrm{ASW}^c$) was normalized as ($\mathrm{ASW}^c$+1)/2. To calculate the batch ASW, we first calculated the batch ASW ($\mathrm{ASW}^b$) for each cell type $j$ as:

$$\mathrm{ASW}_j^b = \frac{1}{|C_j|}\sum_{i \in C_j} 1 - |\mathrm{s}(i)|$$

$$\mathrm{ASW}^b = \frac{1}{|M|}\sum_{j \in \mathrm{M}} \mathrm{ASW}_j^b$$

Here, s($i$) is the silhouette width on batch labels for cell $i$, $C_j$ represents the set of cells with cell labels $j$, and $M$ is the set of unique cell labels.

##### 4.1.5 ARI

The Adjusted Rand Index (ARI) is a measure of similarity between two data clusterings, corrected by chance. An ARI score of 1 represents perfect correspondence, while a value of 0 represents random labeling. To calculate ARI, we performed Louvain clustering using igraph[46] (v.1.3.4) to obtain cluster labels that were compared to the cell type labels. We followed the approach proposed previously[42], where the ARI was optimized based on iterating the clustering resolution from 0.1 to 2 in steps of 0.1. The final clustering was chosen based on the highest ARI value. To compute ARI, we used the function 'ARI' from the 'aricode' R package[47] (v.1.0.0).

##### 4.1.6 NMI

Like ARI, the normalized mutual information (NMI) is a measure of similarity between two data clusterings. An NMI score of 1 represents perfect correspondence, while a value of 0 represents random labeling. To calculate the NMI, we used the optimal clustering obtained from the optimization of the ARI score. To compute NMI, we used the function 'NMI' from the aricode package (v.1.0.0).

#### 5 References and URLs

1. Ding, J. et al. Systematic comparison of single-cell and single-nucleus RNA-sequencing methods. *Nat Biotechnol* **38**, 737-746 (2020).
2. https://singlecell.broadinstitute.org/single_cell/study/SCP424/single-cell-comparison-pbmc-data#study-summary
3. https://github.com/satijalab/seurat-data
4. Segerstolpe, A. et al. Single-Cell Transcriptome Profiling of Human Pancreatic Islets in Health and Type 2 Diabetes. *Cell Metab* **24**, 593-607 (2016).
5. Baron, M. et al. A Single-Cell Transcriptomic Map of the Human and Mouse Pancreas Reveals Inter- and Intra-cell Population Structure. *Cell Syst* **3**, 346-360 e344 (2016).
6. https://bioconductor.org/packages/release/bioc/html/batchelor.html
7. Haghverdi, L., Lun, A.T.L., Morgan, M.D. & Marioni, J.C. Batch effects in single-cell RNA-sequencing data are corrected by matching mutual nearest neighbors. *Nat Biotechnol* **36**, 421-427 (2018).
8. Tabula Muris, C. et al. Single-cell transcriptomics of 20 mouse organs creates a Tabula Muris. *Nature* **562**, 367-372 (2018).
9. https://figshare.com/articles/dataset/Processed_files_to_use_with_scanpy_/8273102
10. Schulte-Schrepping, J. et al. Severe COVID-19 Is Marked by a Dysregulated Myeloid Cell Compartment. *Cell* **182**, 1419-1440 e1423 (2020).
11. https://beta.fastgenomics.org/datasets/detail-dataset-952687f71ef34322a850553c4a24e82e#Files
12. https://beta.fastgenomics.org/datasets/detail-dataset-7ae02f5553074bda92c14a8f0bce2d24#Files
13. Tasic, B. et al. Adult mouse cortical cell taxonomy revealed by single cell transcriptomics. *Nat Neurosci* **19**, 335-346 (2016).
14. Tasic, B. et al. Shared and distinct transcriptomic cell types across neocortical areas. *Nature* **563**, 72-78 (2018).
15. Feng, H. et al. Complexity and graded regulation of neuronal cell-type-specific alternative splicing revealed by single-cell RNA sequencing. *Proc Natl Acad Sci U S A* **118** (2021).
16. https://zhanglab.c2b2.columbia.edu/index.php/Quantas_Documentation
17. https://zhanglab.c2b2.columbia.edu/index.php/OLego
18. Wu, J., Anczukow, O., Krainer, A.R., Zhang, M.Q. & Zhang, C. OLego: fast and sensitive mapping of spliced mRNA-Seq reads using small seeds. *Nucleic Acids Res* **41**, 5149-5163 (2013).
19. Faure, L. et al. Single cell RNA sequencing identifies early diversity of sensory neurons forming via bi-potential intermediates. *Nat Commun* **11**, 4175 (2020).
20. https://github.com/LouisFaure/sensoryfates_paper
21. La Manno, G. et al. RNA velocity of single cells. *Nature* **560**, 494-498 (2018).
22. http://pklab.med.harvard.edu/velocyto/hgForebrainGlut/
23. https://cran.r-project.org/package=Seurat
24. Hao, Y. et al. Integrated analysis of multimodal single-cell data. *Cell* **184**, 3573-3587 e3529 (2021).
25. https://satijalab.org/seurat/
26. Welch, J.D. et al. Single-Cell Multi-omic Integration Compares and Contrasts Features of Brain Cell Identity. *Cell* **177**, 1873-1887 e1817 (2019).
27. https://github.com/welch-lab/liger
28. https://cran.r-project.org/package=harmony
29. Korsunsky, I. et al. Fast, sensitive and accurate integration of single-cell data with Harmony. *Nat Methods* **16**, 1289-1296 (2019).
30. https://github.com/immunogenomics/harmony
31. Polanski, K. et al. BBKNN: fast batch alignment of single cell transcriptomes. *Bioinformatics* **36**, 964-965 (2020).
32. https://github.com/Teichlab/bbknn
33. https://scanpy.readthedocs.io/
34. He, Z., Brazovskaja, A., Ebert, S., Camp, J.G. & Treutlein, B. CSS: cluster similarity spectrum integration of single-cell genomics data. *Genome Biol* **21**, 224 (2020).
35. https://github.com/quadbio/simspec
36. https://cran.r-project.org/package=rsvd
37. McInnes, L., Healy, J. & Melville, J. Umap: Uniform manifold approximation and projection for dimension reduction. *arXiv preprint arXiv:1802.03426* (2018).

38. Moon, K.R. et al. Visualizing structure and transitions in high-dimensional biological data. *Nat Biotechnol* **37**, 1482-1492 (2019).
39. https://cran.r-project.org/package=uwot
40. Butler, A., Hoffman, P., Smibert, P., Papalexi, E. & Satija, R. Integrating single-cell transcriptomic data across different conditions, technologies, and species. *Nat Biotechnol* **36**, 411-420 (2018).
41. https://github.com/immunogenomics/LISI
42. Luecken, M.D. et al. Benchmarking atlas-level data integration in single-cell genomics. *Nat Methods* **19**, 41-50 (2022).
43. Buttner, M., Miao, Z., Wolf, F.A., Teichmann, S.A. & Theis, F.J. A test metric for assessing single-cell RNA-seq batch correction. *Nat Methods* **16**, 43-49 (2019).
44. https://github.com/theislab/kBET
45. https://cran.r-project.org/package=cluster
46. https://cran.r-project.org/package=igraph
47. https://cran.r-project.org/package=aricode
